# Suppression of autoimmune arthritis and neuroinflammation via an amino acid-conjugated butyrate prodrug with enhanced oral bioavailability

**DOI:** 10.1101/2023.04.28.538720

**Authors:** Shijie Cao, Erica Budina, Michal M. Raczy, Ani Solanki, Mindy Nguyen, Taryn N. Beckman, Joseph W. Reda, Kevin Hultgren, Phillip Ang, Anna J. Slezak, Lauren A. Hesser, Aaron T. Alpar, Kirsten C. Refvik, Lucas S. Shores, Ishita Pillai, Rachel P. Wallace, Arjun Dhar, Elyse A. Watkins, Jeffrey A. Hubbell

## Abstract

Butyrate, a metabolite produced by commensal bacteria, has been intensively studied for its immunomodulatory effects on various immune cells, including T regulatory cells, macrophages, and dendritic cells. Butyrate’s development as a drug has been limited by its poor oral bioavailability due to its rapid metabolism in the gut, its low potency and thus high dosing, and its foul smell and taste. By simply esterifying butyrate to serine (*O*-butyryl*-L*-serine, SerBut), a design based on the concept of utilizing amino acid transporters to escape the gut and enhance systemic uptake thus increasing bioavailability, we developed an odorless and tasteless compound for oral administration. In the collagen antibody-induced arthritis (CAIA) and experimental autoimmune encephalomyelitis (EAE) murine models of rheumatoid arthritis and multiple sclerosis, we demonstrated that SerBut significantly ameliorated disease severity, modulated key immune cell populations both systemically and in disease-associated tissues, and reduced inflammatory responses without compromising global immune response to vaccination. Our findings highlight SerBut as a promising next-generation therapeutic agent for autoimmune and inflammatory diseases.

## Introduction

The gut microbiome has been associated with numerous diseases, one of the key mechanisms involving immune regulation through the production of microbial metabolites^1–4^. Among these metabolites, short-chain fatty acids (SCFAs), such as butyrate, have gained significant attention due to their anti-inflammatory and immunomodulatory properties^5–7^. Derived from the microbial fermentation of dietary fiber in the colon, butyrate serves as a primary energy source for colonocytes and maintains intestinal homeostasis^5,8^. It is essential for protecting intestinal barrier function by facilitating tight junction assembly^9,10^. As an epigenetic modulator, butyrate is a histone deacetylase (HDAC) inhibitor and can thus alter chromatin structures and regulate gene expression^11–13^. Through HDAC inhibition, butyrate has been shown to upregulate forkhead box P3 (Foxp3) — a transcription factor involved in the development and function of regulatory T cells (Tregs) — as well as suppress nuclear factor κB (NFκB) activation, inhibit the production of interferon γ (IFNγ), and upregulate peroxisome proliferator-activated receptor γ (PPARγ)^14–17^. In addition to its broad anti-inflammatory activity, butyrate affects immune cell migration, adhesion, cytokine expression, proliferation, activation, and apoptosis^18^. Apart from HDAC inhibition, butyrate can also exert anti-inflammatory effects on immune cells, such as dendritic cells and Tregs, via signaling through specific G-protein coupled receptors (GCPRs): GPR41, GPR43, and GPR109A^19–22^. Collectively, these properties of butyrate hold significant potential for the development of therapeutic strategies, particularly in the treatment of immunological disorders, including autoimmune diseases.

Autoimmune diseases, affecting nearly 5% of the global population, have increased in prevalence over the last few decades^23^. These disorders arise when the immune system mistakenly attacks the body’s own cells and tissues, resulting in chronic inflammation and tissue damage. The onset of autoimmune diseases may be influenced by a combination of genetic and environmental factors. Recent studies have underscored the pivotal role of the gut microbiome in modulating immune responses and influencing the development and progression of autoimmune diseases^3^. For instance, dysregulation of the gut microbiome, or dysbiosis, has been implicated in the pathogenesis of rheumatoid arthritis and multiple sclerosis^24–27^. Current therapeutics for autoimmune diseases, such as immunosuppressive agents, can provide symptom relief but often do not address underlying causes of these complex disorders. There is accumulating evidence that microbial metabolites, such as SCFAs, can impact the immune system and contribute to the development or regulation of autoimmune diseases^28^. Consequently, these findings highlight the potential of using microbial metabolites as therapeutic agents to treat autoimmune diseases by targeting the underlying mechanisms and modulating the immune response.

In spite of this promise, oral administration of sodium butyrate has faced challenges due to its foul, persistent odor and taste, even when administered with enteric coatings or encapsulation. Moreover, butyrate is not absorbed in the gut regions where it could exert therapeutic effects and is rapidly metabolized in the gut as an energy source, limiting its pharmacological impact^29^. Alternative routes of butyrate administration, such as intrarectal delivery or continuous intravenous infusion, are often deemed unfeasible for patients with chronic disorders^30–33^. Therefore, there is a need for innovative delivery methods for butyrate, including prodrugs that can enhance its systemic bioavailability.

To overcome these limitations, we sought to develop a prodrug strategy that enables butyrate to bypass metabolism in the gut, enter the bloodstream, and exert its therapeutic effects systemically after liberation. In this study, we designed an L-serine conjugate of butyrate (*O*-butyryl*-L*-serine or here SerBut) that exploits the gut transport mechanisms of amino acids. Additionally, the conjugation effectively masked the odor and taste of free butyrate, important for facilitating patient compliance for potential clinical applications. We found that SerBut not only enhanced systemic bioavailability, but also facilitated its crossing of the blood-brain barrier (BBB), thereby enabling access to the central nervous system (CNS). In a mouse model of collagen antibody-induced arthritis (CAIA), SerBut treatment showed a significant reduction in disease progression that was associated with a systemic increase in Tregs and an increase in the ratio of immunoregulatory M2 macrophages to pro-inflammatory M1 macrophages. In an experimental autoimmune encephalomyelitis (EAE) model of multiple sclerosis, SerBut significantly prevented disease development and severity, decreased immune cell infiltration in the spinal cord, and upregulated inhibitory markers such as PD-1 and CTLA-4 on CD4^+^ T cells, increased Tregs, and downregulated activation markers on a variety of myeloid cells in the spinal cord-draining lymph nodes. SerBut treatment did not reduce vaccinal immune responses at either the humoral or cellular levels. Thus, in two autoimmune disease models, SerBut administration modulated immune responses at both the myeloid and lymphoid levels without unduly blunting protective immune responses.

## Results

### Conjugation of L-serine to butyrate maintained its biological activity while enhancing oral bioavailability

We synthesized *O*-butyryl*-L*-serine (SerBut) by conjugating L-serine to butyryl chloride using trifluoroacetic acid at room temperature, achieving a 79% yield (**Fig. 1a, fig. S1**). The conjugation effectively masked the unpleasant odor associated with free sodium butyrate or butyrate acid. To assess the HDAC inhibitory activity of SerBut, a property well-recognized in butyrate^19,34^, we performed an *in vitro* histone acetylation assay on the Raw 264.7 macrophage cell line. Our results indicate that SerBut retains HDAC inhibitory activity; however, HDAC inhibition somewhat reduced compared to sodium butyrate (NaBut) at equivalent concentrations (**Fig. 1b**). We propose that during the 18-hour incubation period, SerBut is subject to enzymatic hydrolysis, sequentially releasing butyrate that may, in turn, exert more substantial HDAC inhibition. This is consistent with our design, that SerBut escape then gut and be subsequently hydrolyzed.

**Fig. 1.**
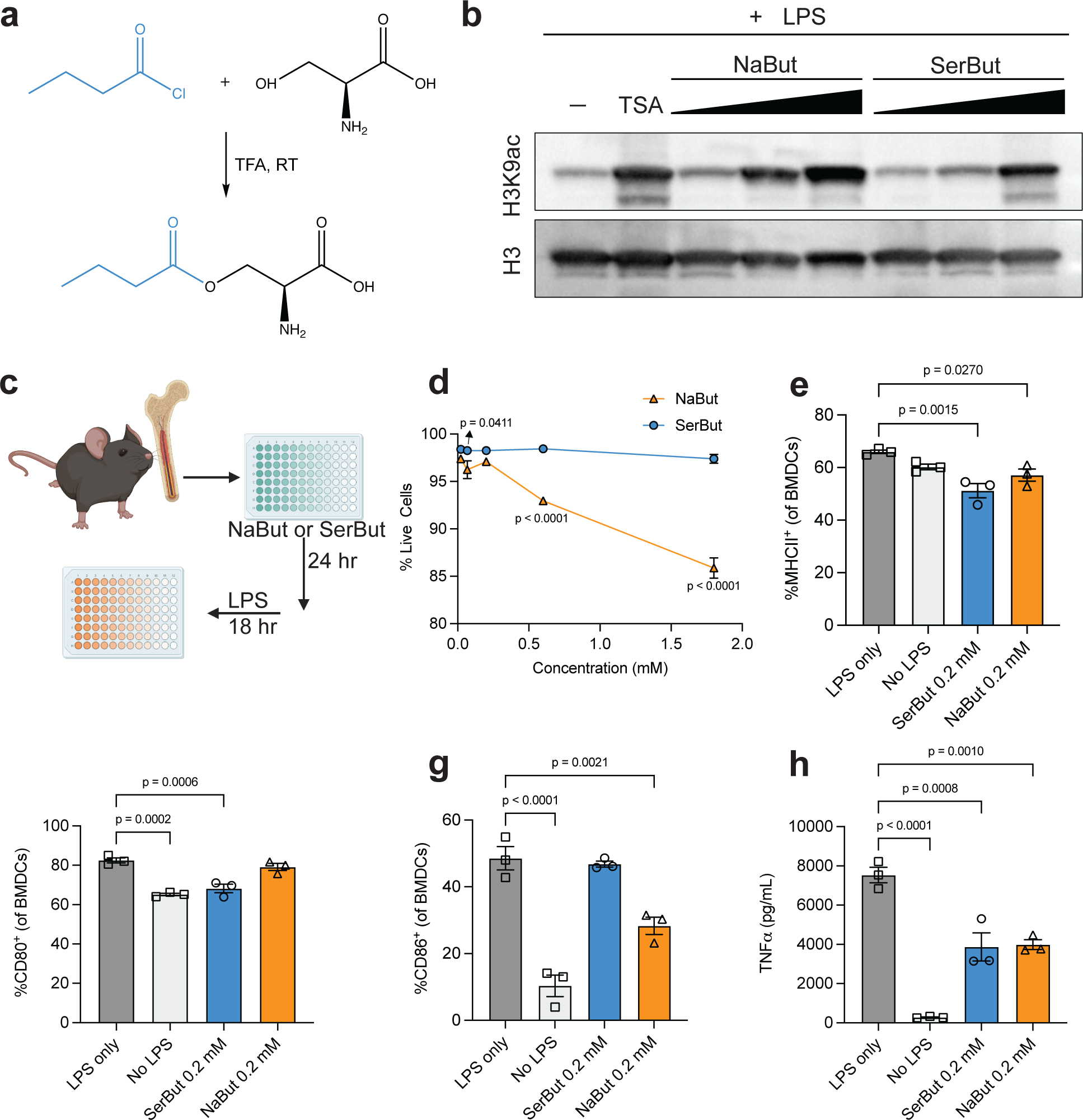
**a.** Chemical synthesis of serine conjugate with butyrate (SerBut). **b.** Whole cell lysates of Raw 264.7 macrophages stimulated with the indicated concentrations of sodium butyrate (NaBut), SerBut, or TSA as well as 100 ng/mL LPS for 18 hr were probed for histone acetylation activity via Western blot. **c.** Experimental schema on bone marrow-derived dendritic cells (BMDCs) incubated with NaBut or SerBut at a series of concentrations for 24 hrs, followed by LPS stimulation for 18 hrs. **d.** Percentage of live BMDCs after treatment. **E-g.** Percentage of CD80^+^, CD86^+^, or MHC class II^+^ BMDCs analyzed by flow cytometry. **h.** TNFα concentration in the cell culture supernatant of BMDCs. *n* = 3, data are representative of two independent experiments. Data represent mean ± s.e.m. Statistical analyses were performed using a one-way ANOVA with Dunnett’s test. P values less than 0.05 were shown. Figure 1c were created with BioRender.com.

Butyrate is known to regulate myeloid cells, including the inhibition of dendritic cell (DC) maturation in response to pro-inflammatory stimuli^35^. We employed bone marrow-derived dendritic cells (BMDCs) to assess the biological effects of SerBut compared to free NaBut (**Fig. 1c**). BMDCs were first incubated with butyrate formulations for 24 hrs, and then stimulated with lipopolysaccharide (LPS) for 18 hrs. We found that NaBut exhibited cytotoxicity to BMDCs at butyrate concentrations above 0.5 mM, while SerBut was well tolerated up to 1.8 mM (**Fig. 1d**). Flow cytometry was used to compare the expression of the BMDC surface markers MHC class II, CD80, and CD86. At the same concentration of 0.2 mM NaBut, SerBut showed similar suppression levels of MHC class II and CD80 (**Fig. 1e, f**). Although SerBut did not suppress CD86 expression as effectively as NaBut at 0.2 mM (**Fig. 1g**), we observed dose-dependent suppression of CD86 with SerBut at higher concentrations (**fig. S2b**). Similarly, SerBut suppressed the secretion of TNFα from BMDCs (**Fig. 1h, fig. S2d**). As demonstrated by the HDAC inhibition assay in Raw 264.7 macrophages and the suppression of LPS-stimulated BMDC activation, SerBut retains the biological activity of butyrate.

The primary site for amino acid absorption is the small intestine, where amino acid transporters are present in intestinal epithelial cells^36^. We conducted a biodistribution study to determine whether conjugating L-serine to butyrate enhances oral butyrate absorption and bioavailability. We measured free butyrate levels in plasma and several major organs after oral gavage of SerBut and compared them with NaBut (**Fig. 2**). We found that SerBut significantly increased plasma butyrate levels at 3 hrs after oral administration compared to NaBut (**Fig. 2b**). In the liver, where orally administered drugs enter directly through the hepatic portal circulation, we observed elevated butyrate levels in mice treated with SerBut but not NaBut (**Fig. 2c**). In the secondary lymphoid organs, we observed elevated butyrate in the mesenteric lymph nodes (mLNs) and spleen (**Fig. 2d, e**). Butyrate levels were also elevated in lungs at 3 hrs post-administration of SerBut (**Fig. 2f**).

**Fig. 2.**
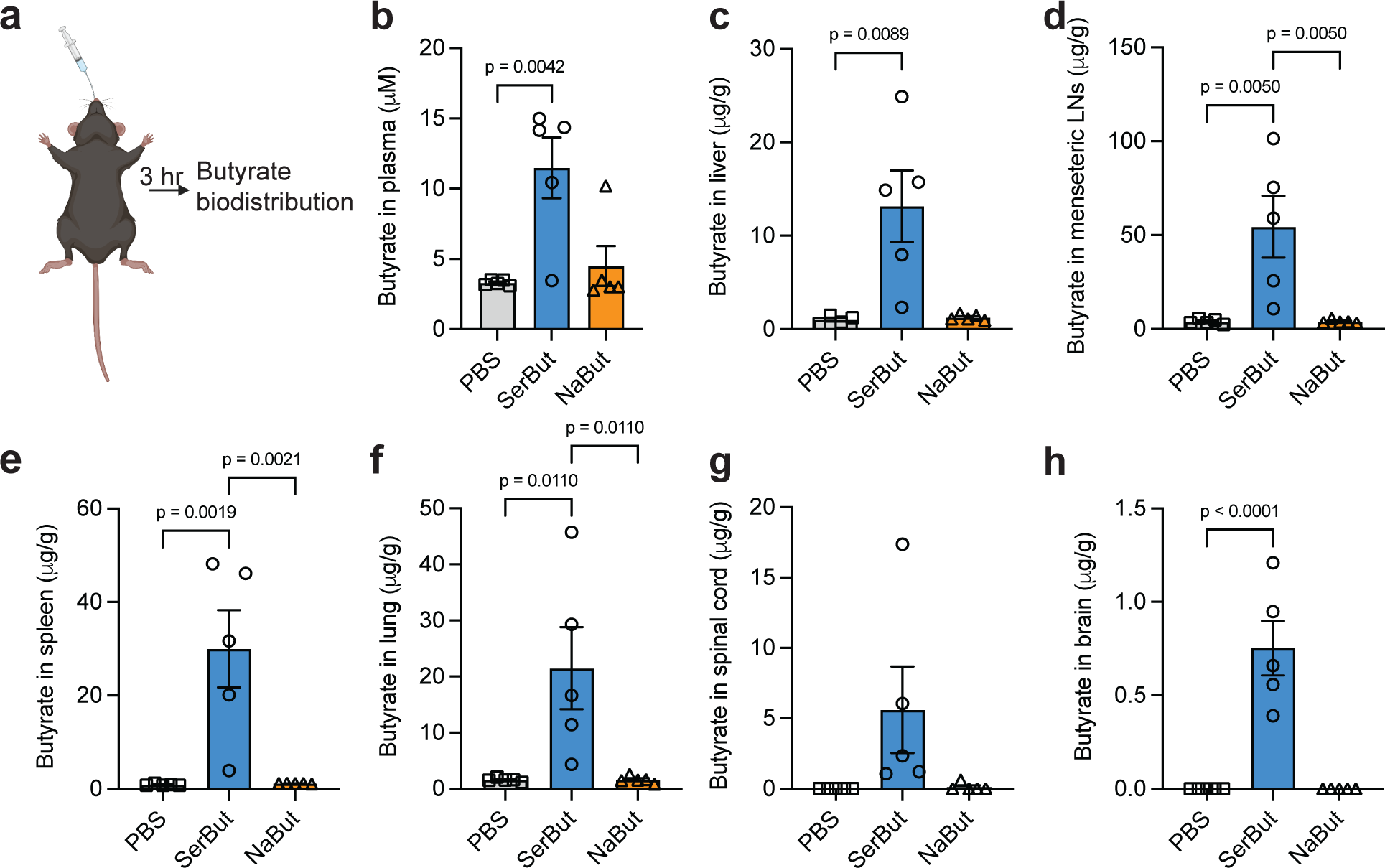
Butyrate biodistribution after SerBut or NaBut oral administration to C57BL/6 mice. The amount of butyrate was detected and quantified in the plasma (**b**), liver (**c**), mesenteric lymph nodes (mLNs) (**d**), spleen (**e**), lung (**f**), spinal cord (**g**), and brain (**h**). Blood was collected by cheek bleeding at 3 hrs post oral gavage. Mice were sacrificed and perfused at 3 hrs post oral gavage, and organs were collected for butyrate quantification. Butyrate was derivatized with 3-nitrophenylhydrazine and quantified by LC-MS/MS. *n* = 5 mice per group. Experiments were repeated twice, data are representative of two independent experiments. Data represent mean ± s.e.m. Statistical analyses were performed using a one-way ANOVA with Dunnett’s test. P values less than 0.05 were shown. Figure 2a were created with BioRender.com.

L-serine is an amino acid known to cross the BBB via the sodium-dependent system A and the sodium-independent alanine, serine, and cysteine transport system^37,38^. Montaser et al. showed that utilizing the L-type amino acid transporter 1 enhances the efficient delivery of small prodrug across the BBB^39^. Therefore, we investigated whether L-serine conjugation could assist butyrate in entering the CNS, including the spinal cord and brain. Our findings revealed that SerBut significantly increased butyrate levels in the spinal cord, reaching approximately 43% of the levels found in the liver when normalized for tissue weight, and in the brain, to about 6%. In contrast, no butyrate was detected in these CNS regions following NaBut administration (**Fig. 2g, h**).

### SerBut suppresses collagen antibody-induced arthritis (CAIA) in mice

The composition of gut microbiota has been shown to affect the development of rheumatoid arthritis (RA)^24,25,40^. Previous research has shown that butyrate can improve RA symptoms by targeting key immune cells such as osteoclasts and T cells^41^. Furthermore, studies have demonstrated that butyrate supplementation effectively reduces arthritis severity in animal models by modulation of regulatory B cells^42^. Given SerBut’s enhanced accumulation in crucial immune tissues following oral administration, we sought to evaluate its efficacy in the CAIA mouse model (**Fig. 3a**). This model is induced by passive immunization with a cocktail of anti-collagen antibodies followed by LPS, which triggers a cascade of innate immune cell infiltration to the joints leading to inflammation and swelling within one week of immunization.

**Fig. 3.**
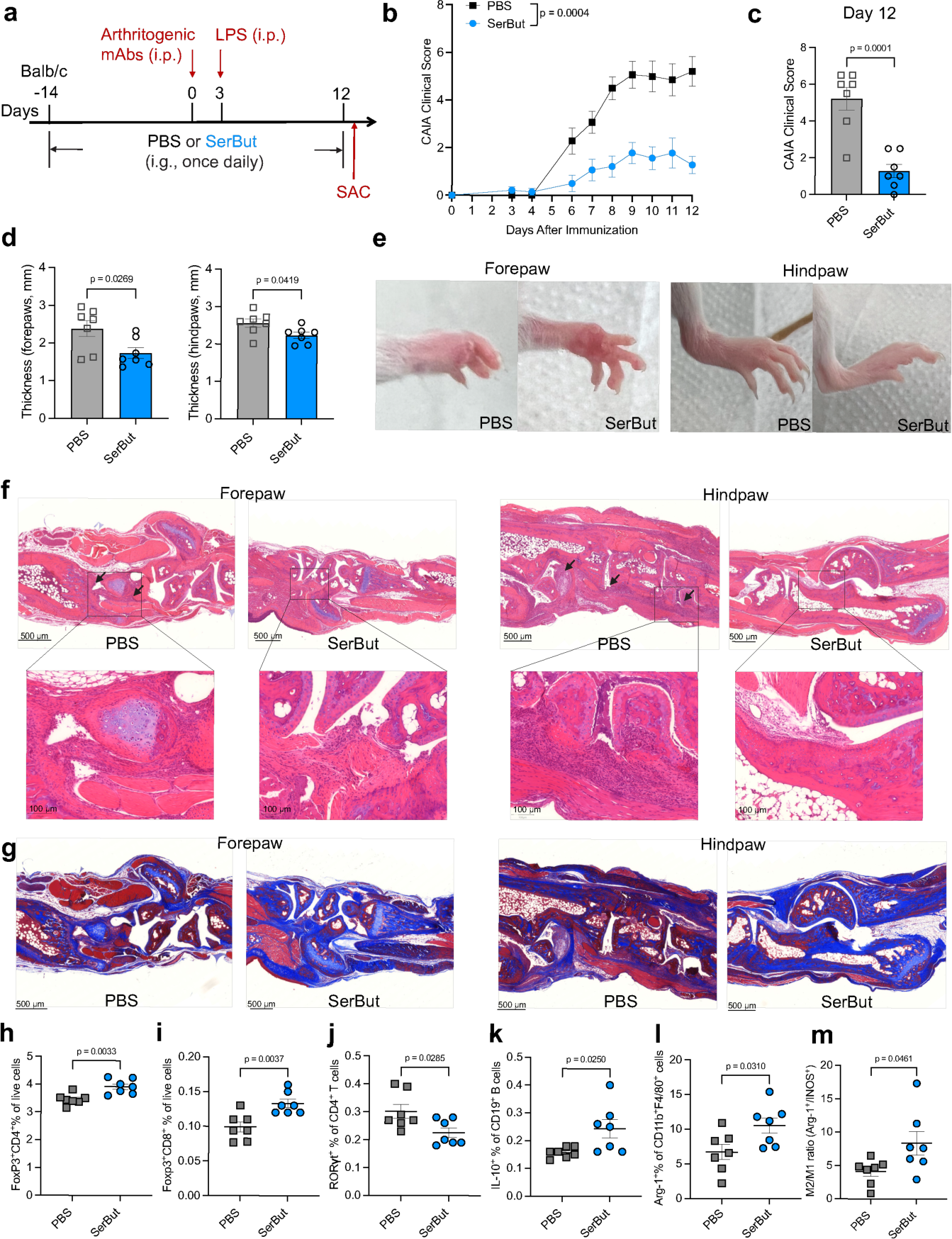
SerBut suppresses arthritis development. **a.** Experimental schema of the collagen antibody-induced arthritis (CAIA) model. Mice were orally gavaged with PBS or SerBut (25 mg) once daily starting on day -14. CAIA was induced by passive immunization with anti-collagen antibody cocktails on day 0, followed by i.p. injection of lipopolysaccharide (LPS). **b.** Arthritis scores in mice measured daily from day 3 after the immunization. **c.** Arthritis scores from PBS or SerBut treated mice on day 12. **d.** The thickness of forepaws or hindpaws measured on day 12 from mice treated with PBS or SerBut. **e.** Representative photos of paws after treatment. **f.** Representative images of mouse joints from paws stained with hematoxylin and eosin on day 12 in each treatment group. Bar = 500 μm (above) or 100 μm (below, in zoomed-in images). Arrows indicate immune cell infiltration. **g**. Representative images of mouse joints from paws stained with Masson’s trichrome on day 12. Bar = 500 μm. Blue represents collagen staining. **h-i.** Percentage of Foxp3^+^ regulatory CD4^+^ T cells (h) or Foxp3^+^ regulatory CD8^+^ T cells (i) of live cells in the spleen measured by flow cytometry. **j-m.** Percentage of RORγt^+^ of CD4^+^ T cells (j) and IL-10^+^ of CD19^+^ B cells (k), as well as Arg-1^+^ of CD11b^+^F4/80^+^ macrophages (l), and M2/M1 macrophage ratio (m) in the hock draining LNs. *n* = 7 mice per group. Experiments were repeated twice, and data are representative of two independent experiments. Data represent mean ± s.e.m. Statistical analyses were performed using Student’s t-test.

Mice were pretreated with SerBut or PBS once daily by oral gavage, beginning two weeks before the induction of arthritis. We observed that SerBut treatment significantly suppressed the development of arthritis in treated mice. In contrast, PBS-treated mice exhibited severe inflammation in the fore- and hindpaws, demonstrated by their increasing clinical scores over time and increasing paw thickness (**Fig. 3b-e**). Histological analysis revealed that oral administration of SerBut effectively suppressed inflammatory responses in the paws and reduced joint pathology compared to PBS-treated mice (**Fig. 3f, g**). In the CAIA model, the anti-collagen antibodies that bind to joint cartilage activate complement proteins, leading to the recruitment of immune cells such as macrophages and T cells to the affected joints^43,44^. This immune cell infiltration contributes to joint inflammation, tissue damage, and the clinical manifestations of arthritis. In our study, we observed that SerBut treatment effectively reduced immune cell infiltration into the joints (**Fig. 3f, fig. S4**). Additionally, anti-collagen antibodies specifically target and degrade collagens in the joints, causing cartilage integrity and joint function loss, ultimately leading to the onset of arthritis symptoms^45^. We found that SerBut treatment also prevented collagen loss in the joints (**Fig. 3g, fig. S5**), suggesting that SerBut may have therapeutic potential in mitigating joint damage and preserving collagen content in the autoimmune arthritis.

To better understand the immunomodulatory effects of SerBut in CAIA, we analyzed immune cell populations in the spleen and hock-draining LNs using flow cytometry. Our results showed that SerBut treatment increased Foxp3 expression in both CD4^+^ and CD8^+^ T cells in the spleen (**Fig. 3h, i**), indicating that SerBut induced expansion of systemic Tregs in the RA disease setting. In the hock-draining LNs (**Fig. 3j**), SerBut significantly reduced the proportion of Th17 cells, as evidenced by the decreased percentage of RORγt^+^ CD4^+^ T cells. Additionally, we observed a significant increase in IL-10-producing B cells within the hock-draining LNs (**Fig. 3k**), which play a vital role in maintaining immune homeostasis^42,46^. Finally, in the hock-draining LNs, SerBut treatment also mediated upregulation of Arg-1 expression in macrophages (**Fig. 3l**), resulting in an increased M2 (Arg1^+^)/M1(iNOS^+^) macrophage ratio in the hock-draining LNs (**Fig. 3m**). The shift towards a higher proportion of immunoregulatory M2-polarized macrophages suggests that SerBut promotes a more balanced immune response, which may be crucial for suppressing inflammation in the paws and reducing joint damage in the context of RA. Overall, these findings highlight the potential therapeutic utility of SerBut in treating RA by modulating the immune system at both the innate and adaptive levels, mitigating disease symptoms.

Next, we compared SerBut to free NaBut in CAIA and investigated whether initiating SerBut treatment post-disease induction (post-administration of the collagen antibody cocktail), with an increased dose regimen, remained efficacious in mitigating disease progression (**Extended Fig. 1**). This approach could be more clinically relevant for rheumatoid arthritis patients who may have circulating autoantibodies for an extended period without exhibiting symptoms^47^. We observed that NaBut did not demonstrate any therapeutic effect. Surprisingly, even during the shortened, 9-day treatment window, SerBut administration proved highly effective in preventing the onset of severe disease. Moreover, our data revealed that this SerBut treatment regimen also promoted the induction of Tregs in both the spleen and hock-draining lymph nodes. These results suggest that Treg induction may play an essential role in the mechanism by which SerBut exerts its immunomodulatory effects in this model.

### SerBut suppresses EAE development via modulation of T cells and myeloid cells in the spinal cord-draining LNs

Multiple sclerosis (MS) is an autoimmune disease in which T cells are reactive to myelin autoantigens, resulting in a chronic demyelinating inflammation in the CNS. Evidence suggests a connection between gut microbiota and MS^26,27,48–52^, particularly dysbiosis in the gut microbiota of MS patients, which includes a reduction of SCFA-producing bacteria^53^. Oral administration of SCFAs, such as butyrate, has been shown to alleviate the severity of EAE, a mouse model of MS, by reducing Th1 cells and increasing Tregs^28,54^. Butyrate has also been demonstrated to ameliorate demyelination and promote remyelination in a mouse model of cuprizone-induced demyelination^55^. Our biodistribution study revealed that SerBut significantly increased butyrate levels in the brain and spinal cord, in addition to the lymphatic tissues, suggesting its potential in treating MS.

We assessed the efficacy of SerBut in suppressing EAE development. To minimize physiological stress from daily oral gavage, mice were administered drinking water containing 100 mM SerBut for the two weeks preceding disease induction and were subsequently administered 24 mg SerBut via oral gavage once daily (**Fig. 4a**). The presence of NaBut or SerBut in the drinking water did not significantly impact the intake of water, as measured in healthy mice (**Fig. S6**). SerBut treatment significantly reduced the EAE clinical score (**Fig. 4b**), indicating disease symptom alleviation. The treatment also delayed EAE onset, as indicated by the lower percentage of SerBut-treated mice with a disease score greater than 1 (**Fig. 4c**). Neither NaBut nor L-serine at equimolar doses prevented disease development.

**Fig. 4.**
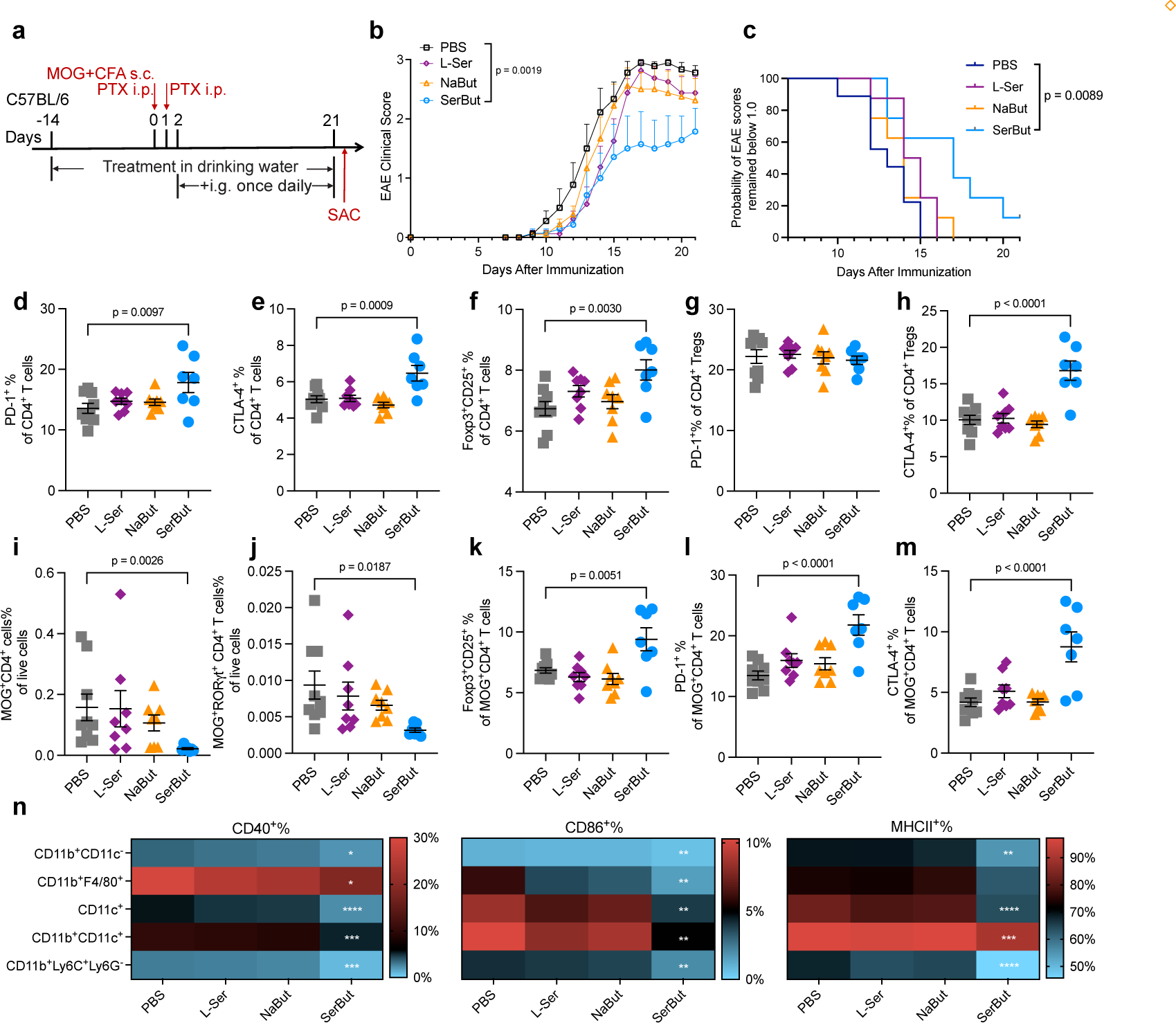
SerBut ameliorates EAE development more effectively than free butyrate or serine. **a.** Experimental schema. EAE was induced in C57BL/6 mice using MOG_35-55_/CFA with pertussis toxin (140 ng). Mice were given drinking water containing 100 mM NaBut, L-Serine (L-Ser), or SerBut from day -14 until the end of the study. On day 2 after EAE induction, PBS (*n* = 9), NaBut (15 mg, molar equivalent to SerBut, *n* = 8), L-Ser (12 mg, molar equivalent to SerBut, *n* = 8), or SerBut (24 mg, *n* = 7) were administered once daily. **b.** Disease progression as indicated by the clinical score. The areas under the curve were compared, and statistical analyses were performed using a one-way ANOVA with Dunnett’s test. **c.** The probability of EAE clinical scores remaining below 1.0 for the three groups. Statistical analysis was performed using the Log-rank (Mantel-Cox) test comparing every two groups. **d-m.** Phenotypes of CD4^+^ T cells from the spinal cord-draining lymph nodes (SC-dLNs, i.e. the iliac and cervical LNs), including the percentage of PD-1^+^ (d), CTLA-4^+^ (e), or Foxp3^+^CD25^+^ (f) of total CD4^+^ T cells; PD-1^+^ (g) or CTLA-4^+^ (h) of Foxp3^+^CD25^+^CD4^+^ Tregs; the percentage of MOG tetramer-positive CD4^+^ (i) or CD4^+^RORγt^+^ (j) T cells of total live cells; and Foxp3^+^CD25^+^ (k), PD-1^+^ (l), CTLA-4^+^ (m) of MOG tetramer-positive CD4^+^ T cells. **n.** Heatmap of the percentage of co-stimulatory molecule (CD40^+^ or CD86^+^) or MHCII^+^ cells of myeloid cells in the SC-dLNs, indicated by the color as shown in the corresponding scale bar. Data represent mean ± s.e.m. Experiments were repeated with similar, though not identical, dosing regimens, and the results were consistent. Statistical analyses were compared between PBS and each treatment group using one-way ANOVA with Dunnett’s test or Kruskal-Wallis test (if not normally distributed). In Figure 4b-m, p values less than 0.05 were shown. In Figure n, *p<0.05, **p<0.01, ***p<0.001, ****p<0.0001. Raw figure and the p-value for Fig. 4n refer to Fig. S9.

A thorough analysis of Immune cell populations in the spinal cord-draining LNs (SC-dLNs) showed that SerBut treatment significantly increased PD-1 and CTLA-4 expression on CD4^+^ T cells, expanded Tregs, and upregulated CTLA-4 expression on CD4^+^ Tregs (**Fig. 4d-h**). Similarly, Treg and PD-1 induction were also observed in CD8^+^ T cells (**fig. S8**). These results suggest that SerBut may help suppress excessive immune responses during EAE by modulating key immune checkpoint markers and regulatory T cell populations. Additionally, in the SC-dLNs, SerBut treatment reduced the number of myelin oligodendrocyte glycoprotein (MOG)-specific T cells and MOG^+^ RORγt^+^ CD4^+^ T cells (Th17 cells), antigen-specific pathogenic cells that contribute to EAE development (**Fig. 4i, j**). SerBut treatment also increased Foxp3, PD-1, and CTLA-4 expression on these MOG^+^ CD4^+^ T cells (**Fig. 4k-m**), potentially preventing these cells from promoting inflammation and tissue damage in EAE. Consistent with the disease readouts, neither NaBut nor L-serine impacted these cellular immune responses.

In the innate immune compartment, SerBut treatment reduced the expression of the co-stimulatory markers, CD40 and CD86, as well as the antigen-presenting molecule MHC class II, on various myeloid cells including CD11b^+^CD11c^-^ cells, CD11b^+^F4/80^+^ macrophages, CD11c^+^ dendritic cells, CD11b^+^CD11c^+^ dendritic cells, and CD11b^+^Ly6C^+^Ly6G^-^ myeloid-derived suppressor cells (MDSCs) (**Fig. 4n**, **Fig. S9**). As these myeloid cells play a crucial role in initiating and propagating immune responses, the reduction of co-stimulatory molecules and MHC class II expression on these cells suggests that SerBut treatment may inhibit their activation and function. This could lead to a dampening of the inflammatory response and contribute to EAE suppression. MDSCs are reported to have immunosuppressive functions, such as inhibiting T cell proliferation and promoting Treg expansion, in the context of EAE^56,57^. We observed not only a downregulation of activation markers on these cells but also a significant increase in the percentage of MDSCs in both SC-dLNs and mesenteric LNs (**Fig. S10**). Overall, our results demonstrate that SerBut has a significant therapeutic impact on EAE by modulating the activity of various immune cell populations and reducing inflammatory responses. All these effects were observed exclusively with SerBut treatment and not with free L-serine nor NaBut, suggesting the importance of conjugation in enhancing butyrate’s systemic and possibly CNS bioavailability.

### SerBut administered post-EAE induction suppresses disease progression and inhibits immune responses in the spinal cord

We next sought to investigate whether administration of SerBut post-EAE induction could also suppress disease progression and induce immunological changes in the spinal cord. Mice were administered twice-daily gavage of SerBut after EAE induction (**Fig. 5a**). We observed that the SerBut-treated mice displayed significantly lower EAE clinical scores compared to PBS-treated mice (**Fig. 5b**). Only 2 out of 8 mice in the SerBut-treated group developed EAE with scores higher than 1 by the end of the study, and the rest remained healthy throughout the experiment (**Fig. 5c**). Additionally, PBS-treated mice experienced a significant drop in body weight starting on day 14, whereas the SerBut-treated mice continued to grow over time (**Fig. 5d**).

**Fig. 5.**
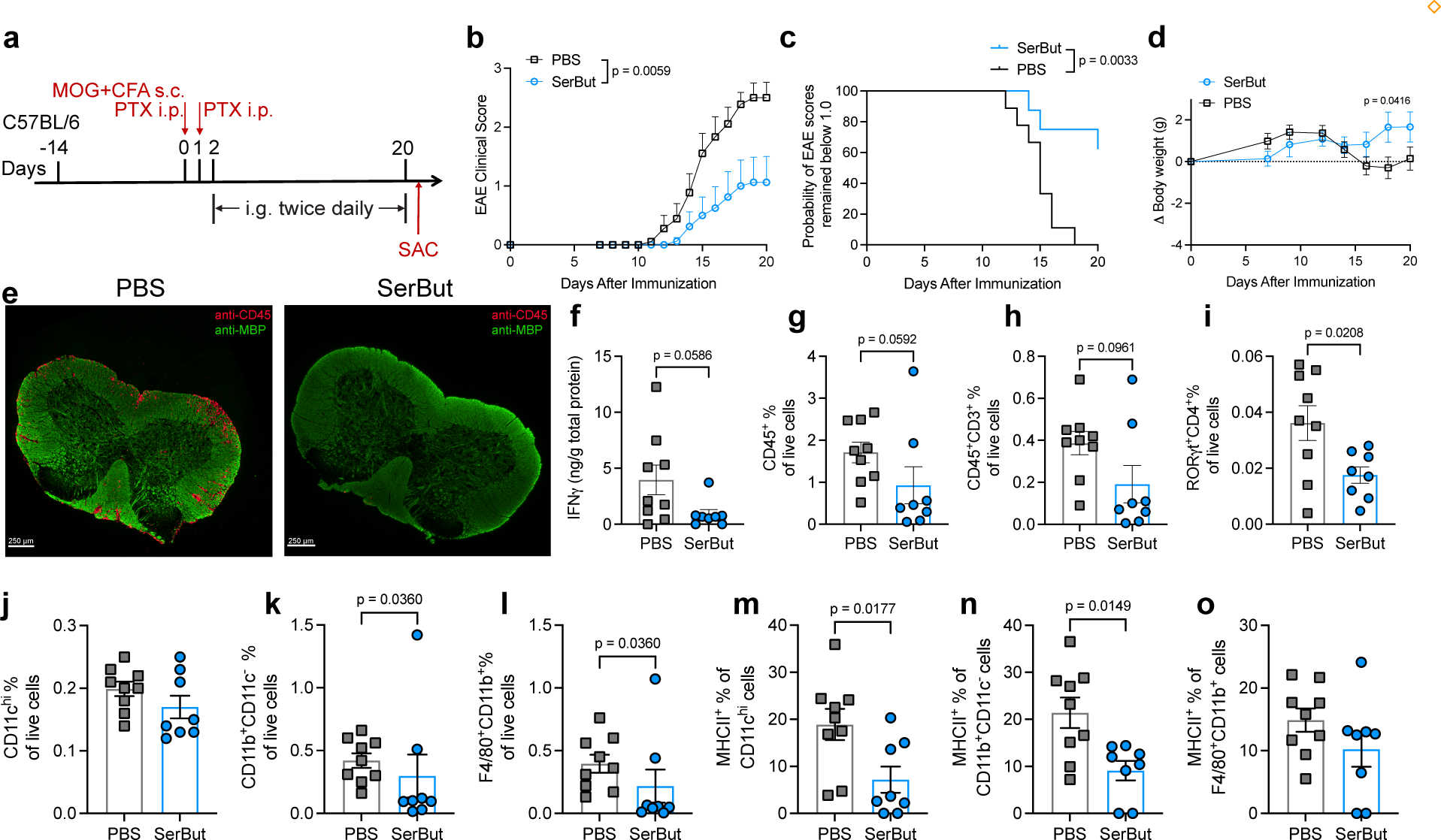
Twice daily gavage of SerBut post-EAE induction suppressed disease progression. **a.** Experimental schema. EAE was induced in C57BL/6 mice using MOG_35-55_/CFA with pertussis toxin (140 ng). On day 2 after EAE induction, PBS (*n* = 9) or SerBut (*n* = 8) at 24 mg/dose were administered twice daily by gavage. **b.** Disease progression as indicated by the clinical score. The areas under the curve were compared, and statistical analyses were performed using Student’s t-test. **c.** The probability of EAE clinical scores remaining below 1.0 in the three groups. Statistical analysis was performed using the Log-rank (Mantel-Cox) test. **d.** Body weight changes. **e.** Representative immunofluorescence images of spinal cord sections from mice treated with PBS or SerBut. Red: anti-CD45 staining; green: anti-myelin basic protein (MBP) staining. Bar = 250 μm. **f.** The concentration of IFN**γ** normalized by the total protein in the spinal cord homogenized supernatant. **g-l.** The percentage of CD45^+^ (g), CD45^+^CD3^+^(h), ROR**γ**t^+^CD4^+^(i), CD11b^+^CD11c^-^(j), F4/80^+^CD11b^+^ (k), and CD11c^hi^(l) of live cells in the spinal cord. **m-o.** The percentage of MHCII^+^ of CD11b^+^CD11c^-^(m), F4/80^+^CD11b^+^ (n), or CD11c^hi^(o) cells in the spinal cord. Data represent mean ± s.e.m. Experiments were repeated with similar, though not identical, dosing regimens, and the results were consistent. Statistical analyses were performed using Student’s t-test.

When PBS-treated mice reached a plateau in disease score, we sacrificed both PBS and SerBut-treated mice and assessed the impact of treatment on the spinal cord. From immunofluorescence images with anti-CD45 and anti-myelin basic protein (MBP) staining, we observed substantial immune cell infiltration into the spinal cords of PBS-treated mice, but not in SerBut-treated mice (**Fig. 5e**, **fig. S12**). The two mice from the SerBut-treated group that did not respond to treatment also showed elevated immune cell infiltration into the spinal cord (**Fig. S12**). Additionally, we homogenized part of the spinal cord and measured IFNγ levels, noticing a decrease in IFNγ in the spinal cords of SerBut-treated mice (**Fig. 5f**).

We also quantified the effect of SerBut treatment on the immune cell compartment in the spinal cord using flow cytometry. We observed a decrease in immune cell populations across most immune cell types in the spinal cord, including CD45^+^ cells, CD3^+^ T cells, and pathological Th17 cells (RORγt^+^ CD4^+^ T cells) (**Fig. 5g-i**), as well as macrophage-like CD11b^+^CD11c^-^ and CD11b^+^F4/80^+^ cells (**Fig. 5j-k**). There was also a notable trend toward a reduction in the infiltration of IL-17A^+^CD4^+^ and IFNγ^+^CD4^+^ cells in the SerBut treated group, although the differences were not significant due to the non-responder mice (**Fig. S13**). Although we did not observe a significant reduction in the CD11c^+^ DC population (**Fig. 5l**), we noted a decrease in MHC class II on their surface, as well as on CD11b^+^CD11c^-^ cells, indicating diminished activation of these cells in the spinal cords of SerBut-treated mice (**Fig. 5m-o**). The reduction of MHC class II on the myeloid cells was observed not only in the spinal cord, but also in the mesenteric LNs (**Extended Fig. 2a, b**). In this experiment, we also observed that SerBut increased the frequency of Foxp3^+^ Tregs both systemically in the spleen and locally in the SC-dLNs and mesenteric LNs (**Extended Fig. 2c**), consistent with our findings in previous RA and EAE experiments. These results suggest that SerBut exerts its immunomodulatory effects both systemically and in disease-related tissues.

We next employed the proteolipid protein (PLP)/CFA-induced EAE relapsing-remitting model to investigate the impact of therapeutic SerBut administration in EAE (**Extended Fig. 3a**). For this purpose, we began treating the mice with PBS or SerBut on day 19 after they had recovered from the first wave of disease. SerBut treatment did not significantly impact relapse onset compared to PBS-treated mice (**Extended Fig. 3b**). This may be attributed to the advanced stage of disease and the relatively short duration of treatment and the. However, with continued SerBut administration, there was a trend towards reduction of relapse compared to PBS-treated counterparts towards the end of the experiment. Intriguingly, flow cytometric analysis of the spinal cord showed a trend towards decreasing immune cell infiltration following SerBut treatment (**Extended Fig. 3c-e**), with a significant upregulation of PD-1 expression on both CD4^+^ and CD8^+^ T cells (**Extended Fig. 3g, Fig. S14**), suggesting that SerBut treatment has the potential to modulate CNS inflammation under therapeutic conditions. While these findings are preliminary, they suggest the potential of SerBut in modulating severe neuroinflammation and merit further exploration of mechanisms of action.

### SerBut does not impact global immune responses to vaccination or alter blood chemistry toxicological markers

Several immune modulators employed in autoimmunity blunt systemic immune responses to vaccination or infection. As a benchmark, we employed fingolimod (FTY720), an FDA-approved oral compound for treating MS^58^. FTY720 targets the sphingosine-1-phosphate receptor and modulates the immune system by sequestering lymphocytes in lymph nodes, reducing their migration to the CNS and ultimately lowering inflammation^59^. Studies have shown that prophylactic oral administration of fingolimod completely prevents the development of EAE in mice, while therapeutic administration reduces the severity of EAE^59^. However, FTY720 treatment can inhibit global immune responses, as demonstrated by our earlier work, where FTY720 administrated to mice vaccinated with ovalbumin (OVA) and adjuvant reduced the generation of OVA-specific IgG^60^.

To gain insight into whether SerBut impacts immune responses to vaccination, we subcutaneously vaccinated naïve mice in the hocks with OVA, in combination with the adjuvants alum and monophosphoryl-lipid A (MPLA), which mimics the clinical vaccine adjuvant AS-04. Mice were treated with either PBS, SerBut, or FTY720 by oral gavage (**Fig. 6a**). Blood collected on days 9 and 13 revealed OVA-specific IgG antibody generation in both PBS and SerBut-treated mice but much less so in FTY720-treated mice, indicating that SerBut does not influence IgG antibody generation, unlike FTY720, in response to OVA vaccination (**Fig. 6b, c**). To evaluate cellular responses, we sacrificed the mice and isolated immune cells from hock-draining LNs. FTY720 reduced B cell (CD19^+^B220^+^), T cell (CD3^+^), and CD4^+^ T cell populations in both hock-draining LNs and the spleen (**Fig. 6d-g, fig. S18a-c**). Interestingly, FTY720 substantially increased Foxp3 Tregs in both LNs and the spleen (**Fig. 6h, i, fig. S18d**). In contrast, SerBut exhibited no impact on any of these cell populations in OVA-vaccinated naïve mice. When we restimulated splenocytes isolated from mice with OVA *in vitro*, we observed a significant reduction in cytokine production, including TNFα, IFNγ, IL-6, IL-5, IL-13, and IL-10 (**Extended Fig. 4**), indicating that FTY720 suppressed antigen-specific T cell responses to OVA. In contrast, SerBut treatment did not suppress these cellular responses, although interestingly, we did observe some significant increases in IL-6, IL-5, IL-13, and IL-10 production with SerBut treatment.

**Fig. 6.**
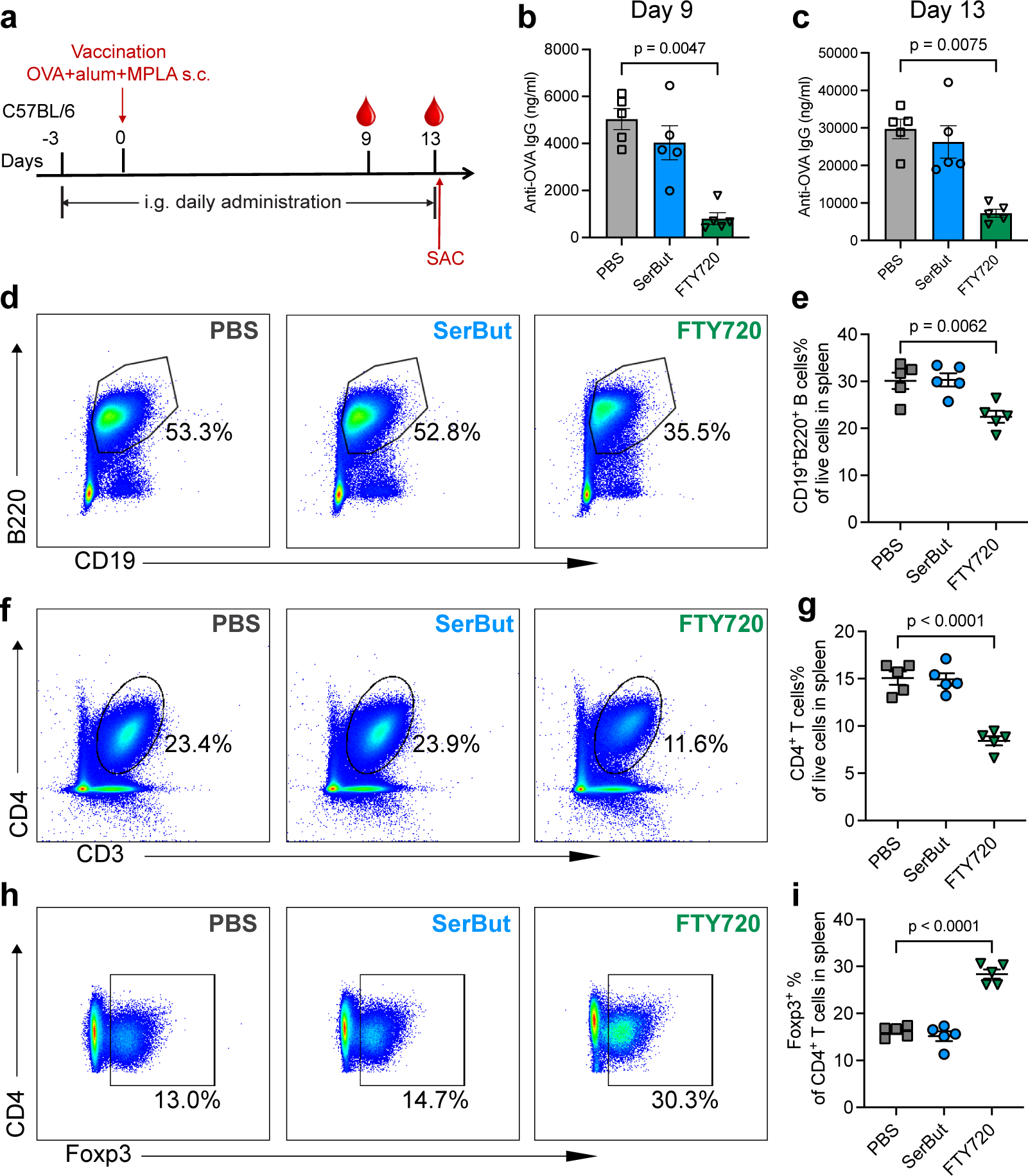
SerBut does not impact immune responses to vaccination compared to FTY720. **a.** Experimental schema. Mice were orally gavaged with PBS, SerBut (twice daily, 25 mg/dose), or FTY720 (once daily, 0.02 mg/dose) starting on day -3 until the end of the experiment. On day 0, mice were immunized subcutaneously in the front hocks with 10 μg endotoxin-free ovalbumin (OVA), 50 μg ALUM, and 5 μg MPLA. **b, c.** Mice were bled on day 9 (b) and day 13 (c), and plasma was analyzed for anti-OVA IgG antibodies. **d-i.** Representative flow cytometry dot plots of CD19^+^B220^+^ (d, e), CD3^+^ (f, g), CD4^+^ (f) and Foxp3^+^CD4^+^ T cells (g) in the spleen, along with their respective percentages of the parental cell population (d, f, h, i), or of the total live cells (e, g). *n* = 5 mice per group. Data represent mean ± s.e.m. Statistical analyses were performed using one-way ANOVA with Dunnett’s post hoc test. P values less than 0.05 were shown.

To determine the impact of SerBut administration on organ function, we also conducted biochemistry analysis on serum from SerBut-treated mice (**Fig. S19**). Overall, we did not observe significant changes in markers of liver, pancreas, and kidney toxicity following SerBut treatment, while FTY720 induced a modest reduction in blood urea nitrogen levels. Taken together, these findings suggest that SerBut does not adversely impact global immune responses to vaccination and is safe for mice when administered via twice-daily oral gavage.

### SerBut induces less pronounced immunological effects on healthy mice than mice with an inflammatory insult

To further investigate the potential impact of SerBut on non-disease bearing mice, we administered PBS, NaBut or SerBut to healthy C57BL/6 mice via twice daily oral gavage and measured the immunological impact on various tissues by flow cytometry. Notably, unlike in autoimmune disease models (CAIA and EAE), SerBut treatment did not elicit CD4^+^ Treg induction in the spleen or lymph nodes of healthy mice (**Extended Fig. 5b**). These results suggest that SerBut may selectively induce Tregs in inflammatory contexts rather than in healthy physiology.

Interestingly, in the spleen, SerBut treatment induced a marked decrease in Th2 cells and an increase in Th17 cells (**Extended Fig. 5c)**. The augmentation of Th17 cells in healthy mice following SerBut administration is particularly intriguing, as it contrasts the reduction of Th17 cells induced by SerBut in the CAIA and EAE models, which are characterized by overabundant, Th17-biased immune responses. This unexpected expansion in Th17 cells warrants further exploration, however it is worth noting that SerBut administration also mediated the expansion of RORγt^+^ Tregs, a distinct population of Tregs known for their enhanced suppressive capacity as compared to Foxp3^+^RORγt^-^ Tregs^61^. Moreover, we observed an upregulation of PD-1 expression on CD4^+^ T cells, whereas CTLA-4 expression remained unchanged (**Extended Fig. 5c)**. These findings in healthy mice suggest differential regulatory pathways may be involved in the modulation of T cell exhaustion markers by SerBut.

In the lamina propria of healthy mice, we also did not observe Treg induction following SerBut treatment (**Fig. S20**). However, SerBut administration reduced the frequency of RORγt^+^ Foxp3^-^ Th17 cells in the lamina propria, presenting an intriguing contrast to its impact on the splenic immune response. These findings prompt further investigation into the disparate Th17 dynamics across tissues. Furthermore, our findings indicate that SerBut’s impact may be more systemic than that of NaBut, which showed no notable changes upon administration in healthy mice. In the mesenteric lymph nodes, SerBut did not alter myeloid cell activation markers, except for a decrease in MHC class II expression on macrophages (**Fig. S21**). This lack of impact on myeloid cells contrasts with observations in the CAIA and EAE models and may be attributed to the fact that cells are less activated during normal physiological conditions. Overall, the impact of SerBut on immune cell populations in healthy mice was minimal and tissue-dependent, particularly regarding Tregs and myeloid cells, where the significant impacts seen in the CAIA and EAE models were absent. The lack of effects in healthy mice suggests a potentially favorable profile for SerBut, as it may reduce the risk of unintended immunosuppression in non-diseased states.

## Discussion

In this study, we investigated the therapeutic potential of SerBut in treating RA and MS using CAIA and EAE as murine models. Orally administered free butyrate can be absorbed and quickly metabolized by host tissues and cells via butyryl-CoA/acetate CoA transferase and phosphotransbutyrylase-butyrate kinase pathways^62^. In the colon, butyrate is primarily consumed by the colonocytes as an energy source^29^. By employing a simple chemical strategy to conjugate serine to butyrate, we improved the oral bioavailability of butyrate including into the CNS, engineering a prodrug candidate that offers several advantages, including a lack of unpleasant odor and taste, and higher efficacy compared to free butyrate. Our results demonstrated that SerBut significantly ameliorated the severity of both diseases, modulated key immune cell populations, and reduced inflammatory responses, all without compromising global immune responses to vaccination.

Butyrate, a key metabolite from commensal bacteria, has multiple modulating effects across different types of immune cells. It has been shown to facilitate peripheral generation of Tregs through both direct upregulation of Foxp3 by HDAC inhibition^16,17^, and indirect effects from the induction of tolerogenic DCs^19,63^. In our study, we demonstrated that SerBut treatment increased peripheral Tregs in both CAIA and EAE settings, as well as in both local draining LNs and spleen. Notably, we observed enhanced biodistribution of SerBut in these tissues after oral administration, which is not limited to gut tissues as previously reported^64,65^. Butyrate has also been shown to modulate the differentiation of macrophages and DCs. Through HDAC inhibition, butyrate downregulates LPS-induced proinflammatory cytokines produced by macrophages^34^, and activates macrophage metabolism towards oxidative phosphorylation through upregulation of Arg-1, promoting an anti-inflammatory M2 phenotype^66^. In our CAIA model, enhancing the systemic bioavailability of butyrate through SerBut significantly increased the M2/M1 macrophage ratio in hock-draining LNs, which could exert direct anti-inflammatory effects on paw and joint inflammation. Moreover, butyrate has been shown to suppress DC activation and induce tolerogenic DCs through a combination of signaling via GPR109A and HDAC inhibition^19^. In our study, we observed that SerBut suppresses LPS-induced activation, including co-stimulatory markers CD80 and CD86, and MHC class II expression of BMDCs isolated from mice compared to NaBut. Consistently, in the EAE model, we observed downregulation of these costimulatory markers and MHC class II on various myeloid cells across different tissues, suggesting the important roles butyrate can play in suppressing disease progression through this pathway while delivered systemically.

Myeloid cells, such as DCs and macrophages, play a crucial role in antigen presentation and T cell activation. For instance, CD86 interacts with CD28 on T cells, promoting their activation and proliferation^67,68^. CTLA-4 is a surface receptor predominantly expressed on T cells, particularly on Tregs, and competes with CD28 for binding to CD86 on myeloid cells, delivering an inhibitory signal to T cells^69^. Intriguingly, in our EAE experiment, we observed that SerBut treatment led to a significant increase in CTLA-4 expression on both total CD4^+^ T cells, and Tregs in the spinal cord-draining LNs. This concurrent upregulation of CTLA-4 on T cells, and downregulation of CD86 on myeloid cells can synergistically contribute to a more profound suppression of immune activation, ultimately dampening autoimmune responses.

In EAE, we have shown that twice-daily administration of SerBut significantly reduces immune cell infiltration into the spinal cord. In future studies, we will investigate whether these effects on the CNS effect are modulated directly by butyrate that crosses the BBB or indirectly due to butyrate’s effects on peripheral immune cells which subsequently induce immunomodulation in the CNS. Additionally, the effect of butyrate on BBB (or blood-spinal cord barrier) endothelial cell functions and barrier integrity warrants further investigation.

Interestingly, we noticed that SerBut’s immunomodulatory effects are context-dependent, promoting immune homeostasis in inflammatory settings. This is evidenced by our findings that SerBut induces Treg induction in the autoimmune disease models such as CAIA and EAE (**Fig. 3h, i, Fig. 4f, k, Extended Fig. 1d-g, Extended Fig. 2c, fig. S8**), but not following administration in healthy mice or OVA-vaccinated mice (**Fig. 6h, Extended Fig. 5b, fig. S18d, fig. S20a, b**). We observed similar context-dependent differences in Th17 responses: SerBut treatment in disease models with Th17-biased autoimmunity induced a reduction in Th17 cells following treatment (**Fig. 3j**, **Fig. 4j**, **Fig. 5i, Extended Fig. 3e**), while in healthy mice, the impact of SerBut administration was more nuanced and tissue-specific (**Extended Fig. 5c, fig. S20c, d**). Moreover, the impact of SerBut on myeloid cells was also more pronounced in the presence of inflammation (**Fig. 3l, m, Fig. 4n**, **Fig. 5m-o, Extended Fig. 2a, b, fig. S9, fig. S10**) than in healthy mice, where myeloid cells were not pre-activated (**Fig. S21**). The differential, disease-dependent immunological effects of SCFAs are corroborated by other literature across various fields, including cancer immunotherapy and vaccine responses^70,71^.

We have shown that SerBut has several potential benefits over free NaBut for clinical translation: serine conjugation to butyrate masks its odor and taste, yields higher bioavailability, and shows greater efficacy in suppressing autoimmune disease models compared to currently available NaBut. The dose used in these studies can be converted to 6 grams of SerBut per dose for human clinical trials, which could be formulated as powder that could be dissolved in drinking water for daily consumption. Importantly, SerBut did not adversely impact global immune responses to vaccination, as demonstrated by the generation of equivalent OVA-specific IgG humoral and cellular immune responses in both PBS and SerBut-treated mice. Although we observed effective immune modulation in the context of autoimmune arthritis and EAE, the immune response elicited by a strong, Th1-biasing adjuvant (the TLR4 agonist MPLA in the alum depot) during vaccination was sufficient to overcome the effects of SerBut^72^. This distinction offers a compelling insight into the potential of SerBut as a more targeted therapeutic strategy for chronic autoimmune diseases, with a possibly lower risk of the adverse effects associated with broad immunosuppression.

Our study provides evidence that SerBut has potential as a next-generation therapeutic agent for RA and MS. Further studies, including preclinical and clinical studies, are needed to better understand the long-term safety and efficacy of SerBut in the context of rheumatoid arthritis, multiple sclerosis, and other autoimmune and inflammatory diseases. Given its broad immunomodulatory effects shown in our study, it would be valuable to explore the potential of SerBut in treating a broader range of immune-related conditions.

## Materials and Methods

### Study design

The objective of this study was to chemically conjugate L-serine to butyrate to improve butyrate’s oral bioavailability and investigate the therapeutic potential of the conjugate, SerBut, in the context of RA and MS using CAIA and EAE murine models.

In the biodistribution study, we quantified the butyrate content in various major organs following oral gavage of SerBut. In the RA model, mice were treated daily with either PBS, NaBut, or SerBut via oral gavage. Paw inflammation was assessed over time, and the pathology of inflamed paws and joints was evaluated using histology. Immune cell phenotypes were analyzed by flow cytometry at the end of the experiment when PBS-treated mice reached a plateau in disease scores. In the EAE model, we compared the efficacy of SerBut, PBS, free butyrate, and free L-serine in preventing disease progression. Immune responses were evaluated at the end of the experiment when PBS-treated mice reached a plateau in disease scores. In the vaccination study, the global immunosuppressive effects of SerBut were compared to those of FTY720. The study endpoint was determined based on previous reports that day 13 after OVA immunization was sufficient to induce anti-OVA IgG antibodies^60^.

Sample size was determined using results obtained from previous and preliminary studies. At least 5, and in most cases 7-9 independent biological replicates were examined for each group analyzed. See figure legends for details on n for each display figure. All experiments were replicated at least twice. Mice were randomly assigned to treatment groups, except in the PLP-EAE study where mice with already-established disease were assigned to treatment groups based on average clinical score. The person assessing clinical scores for RA and EAE experiments was separate from the person administering treatment and was blinded to the treatment group. The person performing fluorescent imaging and histology analysis was also blinded to treatment groups. Statistical methods are described in the “Statistical analysis” section.

### Synthesis of SerBut

L-Serine (20 g, 0.19 mol) was added to trifluoroacetic acid (200 ml) and the suspension was stirred for 30 min until everything dissolved. Butyryl chloride (25.7 ml, 0.23 mol) was then added to the solution and the mixture was stirred for 2 hrs at room temperature. The reaction was then transferred to an ice bath and diethyl ether (500 ml) was added, which resulted in a precipitation of a white solid. The resultant fine white precipitate was collected by filtration, washed with cold diethyl ether, and dried under vacuum to afford 26.3 g of *O*-butyryl*-L*-serine (0.15 mol, 79%). The final product was confirmed by 1H NMR (500MHz, DMSO-d6) [ppm]: 0.88 (3H, t), 1.55 (2H, m), 2.32 (2H, t), 4.30 (1H, t), 4.43 (2H, d), 8.66 (2H, s), 14.06 (1H, s).

### Mice

C57BL/6 mice, aged 8-12 weeks, were purchased from Charles River (strain code: 027, Charles River). BALB/c mice, aged 6-10 weeks were purchased from the Jackson Laboratory (strain code: 000651, JAX). SJL/JCrHsd mice, aged 6 weeks, were purchased from Envigo (strain code: 052, Envigo). C57BL/6, BALB/c, and SJL/JCrdHsd mice were maintained in a specific pathogen-free (SPF) facility at the University of Chicago. Mice were maintained on a 12 hr light/dark cycle at a room temperature of 20-24 °C. All protocols used in this study were approved by the Institutional Animal Care and Use Committee of the University of Chicago.

### Flow cytometry and antibodies

Flow cytometry was performed using BD LSRFortessa^TM^, and data were analyzed using FlowJo version 10.8.0. Antibodies against the following markers were used in the RA and EAE mouse models: INOS (BUV737, Cat#367-5920-82, Invitrogen), CD11c (BV421, Cat#562782, BD Biosciences), CD11c (BV785, Cat#563735, Biolegend), Ly6C (BV605, Cat#128036, BioLegend), CD11b (BV650, Cat#563402, BD Biosciences), CD11b (BV711, Cat#101242, BioLegend), Ly6G (AF488, Cat#127626, BioLegend), CD40 (PerCP/Cy5.5, Cat#124624, BioLegend), CD40 (BUV615, Cat#751646, BD Biosciences), CD206 (PE, Cat#141706, BioLegend), CD206 (AF700, Cat#141734, Biolegend), Arginase 1 (PE-Cy7, Cat#25-3697-82, Invitrogen), F4/80 (APC, Cat#123116, BioLegend), F4/80 (PE, Cat#565410, BD Biosciences), CD86 (AF700, Cat#105024, BioLegend), CD86 (BUV395, Cat#564199, BD Biosciences), I-A/I-E (APC/Cy7, Cat#107628, BioLegend), I-A/I-E (BV421, Cat#107632, Biolegend), CD19 (BUV396, Cat#563557, BD Biosciences), CD3 (BUV737, Cat#741788, BD Biosciences), CD3 (BV605, Cat#100351, BioLegend), CD3 (APC-Fire750, Cat#100362, Biolegend), CD4 (BV605, Cat#100548, BioLegend), CD4 (BV711, Cat#100550, BioLegend), CD4 (BUV496, Cat#612952, BD Biosciences), CD4 (AF647, Cat#553051, BD Biosciences), PD-1 (BV711, Cat#135231, BioLegend), PD-1 (APC-Cy7, Cat#, BD Biosciences), Foxp3 (AF488, Cat#53-5773-82, Invitrogen), PD-L1 (BV711, Cat#563369, BD Biosciences), RORγt (PerCP/Cy5.5, Cat#562683, BD Biosciences), RORγt (APC, Cat#562682, BD Biosciences), RORγt (BV421, Cat#562894, BD Biosciences), CD5 (PE, Cat#100607, BioLegend), CTLA-4 (PE-Cy7, Cat#25-1522-80, Invitrogen), CTLA-4 (PE-Cy7, Cat#106314, Biolegend), CD25 (APC, Cat#162105, BioLegend), CD25 (PerCP/Cy5.5, Cat#561112, BD Biosciences), CD25 (BV650, Cat#10238, Biolegend), CD8 (AF700, Cat#100730, BioLegend), CD8 (BUV737, Cat#612759, BD Biosciences), IL-10 (APC/Cy7, Cat#505036, BioLegend), CD45 (BUV395, Cat#564279, BD Biosciences), CD45 (V450, Cat#560501, BD Biosciences), and CD45 (BUV805, Cat#748370, BD Biosciences). The I-A(b) mouse MOG 38-49 GWYRSPFSRVVH (MOG tetramer, PE) was obtained from NIH Tetramer Core Facility.

### *In Vitro* Histone Acetylation assay

Raw 264.7 macrophages were cultured to 50% confluency before stimulation with the indicated concentration of inhibitor with 100ng/mL LPS (Invivogen tlrl-smlps) and cultured for 18hrs. After stimulation, cells were harvested via scraping before histone isolation according to manufacturer’s protocol (abcam 113476). Protein gels were then loaded with 8 μg of total protein isolate as quantified via BCA (Pierce 23225) before semi-dry transfer. The membrane was blocked with 5% BSA in PBS-T before probing with anti-total H3 (CST 4499) and anti-acetylated H3K9 (CST 9649) at 1:1000 in 2% BSA in PBST, followed by the HRP-conjugated secondary antibody (Southern Biotech 4050-05) at 1:10,000 in 2% BSA in PBS-T and detection.

### Mouse BMDC isolation and activation study

Murine BMDCs were collected from 6-week-old female C57BL/6 mice as described by Lutz et al^73^. BMDCs were seeded at 3 x 10^6^ total cells/plate in petri dishes. Cells were cultured at 37°C and 5% CO_2_ in the media: RPMI 1640 (Life Technologies), 10% HIFBS (Gibco), GM-CSF (20 ng/mL; recombinant Mouse GM-CSF (carrier-free) from BioLegend), 2 mM l-glutamine (Life Technologies), 1% antibiotic-antimycotic (Life Technologies). Media was replenished on day 3 and day 6. Cells were used on day 9. Isolated BMDCs were plated in round-bottom 96 well plates at 100,000 cells per well in RPMI media and co-cultured with different concentrations of either NaBut or SerBut (from 0.02 to 1.8 mM) for 24 hrs. Subsequent to addition of butyrate compounds, cells were stimulated with LPS (1 μg/mL) for another 18 hrs. The supernatant of cell culture was collected and analyzed by LEGENDPlex^TM^ to analyze the concentrations of cytokines (BioLegend). BMDCs were collected and stained with live/dead dye (Cat#L34957, Invitrogen) and fluorescent antibodies against CD11c (PE-Cy7, Cat#558079, BD Biosciences), MHC class II (APC-Cy7, Cat#107628, BioLegend), CD80 (PE, Cat#104708, BioLegend), CD86 (FITC, Cat#MA1-10300, Invitrogen). Cell phenotype was analyzed using flow cytometry (BD LSRFortessa).

### Biodistribution of SerBut

C57BL/6 mice were orally administered with 50.4 mg NaBut or 80 mg SerBut (both containing equivalent 40 mg butyrate). At 3 hrs post-administration, mice were anesthetized under isoflurane and blood was collected via cheek bleeding, and mice were then transcardially perfused with a minimum of 30 mL PBS containing 1 mM EDTA. Organs, including liver, mesenteric lymph nodes (mLNs), spleen, lung, spinal cord, and brain were collected, immediately frozen in dry ice, and then transferred to -80°C until further processing.

To extract butyrate from plasma or organs, a 1:1 v/v acetonitrile (ACN) to water solution was used. Plasma was mixed 1:1 with the ACN/water solution and centrifuged to remove denatured proteins. Organs were weighed, transferred to Lysing Matrix D tubes, and combined with the 1:1 v/v ACN/water solution. Samples were then lysed using a FastPrep-24 5G homogenizer (MP Biomedicals) and centrifuged. The supernatants were collected for butyrate measurement.

Samples were prepared and derivatized as described previously^7,74^. A 3-nitrophenylhydrazine (NPH) stock solution was prepared at 0.02 M in water:ACN 1:1 v/v. A 1-ethyl-3-(3-dimethylaminopropyl)carbodiimide (EDC) stock solution (with 1% pyridine added) was prepared at 0.25 M in water:ACN 1:1 v/v. The internal standard, 4-methylvaleric acid, was added. Samples were mixed with NPH and EDC stocks at a 1:1:1 volume ratio. The mixture was heated in a heating block at 60°C for 30 min. Samples were then filtered through 0.22 μm filters and transferred into HPLC vials, which were stored at 4°C before analysis.

An Agilent 6460 Triple Quad MS-MS was used to detect the derivatized butyrate. Both derivatized butyrate-NPH and 4-methylvaleric-NPH were detected in negative mode. Column: ThermoScientific C18 4.6 × 50 mm, 1.8 μm particle size, at room temperature. Mobile phase A: water with 0.1% v/v formic acid. Mobile phase B: acetonitrile with 0.1% v/v formic acid. Injection volume: 5.0 μL. Flow rate: 0.5 mL/min. Gradient of solvent: 15% mobile phase B at 0.0 min; 100% mobile phase B at 3.5 min; 100% mobile phase B at 6.0 min; 15% mobile phase B at 6.5 min. The MS conditions were optimized using pure butyrate-NPH or 4-methylvaleric-NPH at 1 mM. The fragment voltage was set to 135 V, and the collision energy was 18 V. Multiple reaction monitoring (MRM) of 222 → 137 was assigned to butyrate, and MRM of 250 → 137 was assigned to 4-methylvaleric acid as the internal standard. The ratio between MRM of butyrate and 4-methylvaleric acid was used to quantify butyrate concentration. The final butyrate content in each organ was normalized by organ weight.

### SerBut Administration in Naïve C57BL/6 mice

C57BL/6 mice, aged 8 weeks, were purchased from Charles River Laboratories and housed in the animal facility at the University of Chicago for 2 weeks before use. From day 0 to day 10, mice were administered twice daily oral gavage of PBS, NaBut (15 mg, molar equivalent to SerBut), or SerBut (24 mg). On day 10, mice were sacrificed. The lymph nodes (cervical and illiac, mesenteric, and hock draining), spleen, and lamina propria were harvested and processed for flow cytometry analysis.

### Collagen-antibody inducing arthritis (CAIA) model

BALB/c mice, aged 6 weeks, were purchased from the Jackson Laboratory and housed in the animal facility at the University of Chicago for 2 weeks before immunization. Mice were rally gavaged with PBS or SerBut (25 mg) once daily starting on day -14 or with PBS, NaBut (15 mg, molar equivalent to SerBut), or SerBut (24 mg) twice daily beginning on day 3 at the age of 8 weeks. CAIA was induced by passive immunization with an anti-collagen antibody cocktail (1 mg per mice by i.p. injections, Arthrogen-CIA® 5-Clone Cocktail Kit, Chondrex, Inc.) (on day 0, followed by an intraperitoneal injection of LPS (25 μg) on day 3. Mice cages were layered with soft (pine) bedding throughout the experiment. Arthritis severity was monitored daily after day 3 using the criteria for clinical scores established by Chondrex, Inc., as described previously^60^.

On day 12, the thickness of mouse fore- and hindpaws was measured to assess the swelling resulting from arthritis. On day 13, mice were sacrificed, and the spleen and hock-draining LNs, including popliteal, axillary, and brachial LNs, were harvested for immunostaining followed by flow cytometry analysis.

The paws were collected for histological analysis, as described previously^60^. Briefly, paws were fixed in 2% paraformaldehyde (Thermo Scientific), decalcified in Decalcifer II (Leica), and stored in 70% ethanol until paraffin embedding. Paraffin-embedded paws were sliced into 5 μm-thick sections and stained with hematoxylin and eosin, or Masson’s trichrome. The images were captured using a CRi Panoramic SCAN 40x or MIDI 20x Whole Slide Scanner, or Olympus VS200 Slideview Research Slide Scanner, and analyzed using ImageJ and QuPath software.

### Experimental autoimmune encephalomyelitis (EAE) model

C57BL/6 female mice (7-8 weeks old) were purchased from Charles River Laboratories and housed in the animal facility at the University of Chicago for 2 weeks before immunization. Female C57BL/6 mice, aged 10 weeks, were subcutaneously immunized at the dorsal flanks with an emulsion of MOG_35-55_ in complete Freund’s adjuvant (MOG_35-55_/CFA Emulsion, Hooke Laboratories) on day 0, followed by i.p. administration of pertussis toxin (140 ng) in PBS on both day 0 and day 1. The development of EAE was monitored, and clinical scores were measured daily from day 7 to day 20. The criteria for clinical scores was according to the instructions from Hooke Laboratories and described previously^75^.

In the experiment from Fig. 3, Mice were given drinking water containing 100 mM NaBut, L-Serine (L-Ser), or SerBut from day -14 until the end of the study. On day 2 after EAE induction, PBS, NaBut (15 mg, molar equivalent to SerBut), L-Ser (12 mg, molar equivalent to SerBut), or SerBut (24 mg) were administered by once daily oral gavage. In the experiment from Fig. 4, mice were administered of PBS or SerBut (24 mg) twice daily by oral gavage from day 2 after EAE induction.

On day 21 or 22, mice were sacrificed. The spinal cords were collected and separated into three sections for immunofluorescence imaging, cytokine measurement through homogenization, or immunostaining for flow cytometry analysis. Blood was collected through cardiac puncture, and spleen, mesenteric LNs, and spinal cord-draining LNs (SC-dLNs, including cervical LNs and iliac LNs) were harvested. Single-cell suspensions were collected for the immunostaining followed by the flow cytometry analysis. Major cytokines from the plasma and spinal cord after homogenization were analyzed via LEGENDPlex (BioLegend).

SJL/JCrHsd female mice (6 weeks old) were purchased from Envigo Laboratories and housed in the animal facility at the University of Chicago for 2 weeks before immunization. Female SJL/JCrHsd mice, aged 8 weeks, were subcutaneously immunized at the dorsal flanks with an emulsion of MOG_35-55_ in complete Freund’s adjuvant (PLP_139-151_(naïve) /CFA Emulsion, Hooke Laboratories) on day 0, followed by i.p. administration of pertussis toxin (100 ng) in PBS on both day 0 and day 2. The development of relapsing-remitting EAE was monitored, and clinical scores were measured daily from day 7 to day 40. Clinical score was determined according to the instructions from Hooke Laboratories and described previously^75,76^. On day 19 after EAE induction, mice were assigned to treatment groups. Mice in each treatment group had the same average clinical score. From day 19 to endpoint, mice were administered twice daily oral gavage of PBS or SerBut (24 mg). On day 40, mice were sacrificed and tissues were processed for flow cytometry analysis as described above.

### Immunofluorescence imaging of spinal cord sections

Thoracic and lumbar spines of EAE mice were collected. The tissues were fixed in 2% paraformaldehyde (Thermo Scientific) and then stored in 70% ethanol until paraffin embedding. Paraffin-embedded spinal cords were sliced into 5 μm-thick sections as previously described^60,75^. The sections were deparaffinized through a series of washes in xylene, ethanol, and double-distilled water. Spinal cord sections were immersed in each solution for 2 min per wash. Antigen retrieval was performed using 1x pH 6.0 citrate buffer at 50-55°C for 45 min. Sections were blocked for 1 hr at room temperature in PBS containing 0.3% Triton-X and 5% normal goat serum. Primary antibodies against CD45 (clone 30-F11, BioLegend) and MBP (clone ab40390, Abcam) were applied at 1:100 dilution in blocking buffer and incubated for 16 hrs at 4°C. Sections were washed and then incubated with donkey anti-rat IgG (H+L) AF647 (A48272, Invitrogen) and donkey anti-rabbit IgG (H+L) AF488 (2340683, Jackson ImmunoResearch) secondary antibodies for 16 hrs at 4°C. Following additional washes, sections were mounted using Fluoromount-G Mounting Medium and imaged with an Olympus IX83 spinning-disc confocal fluorescence microscope. Image processing was performed using ImageJ and QuPath software.

### Evaluation of immune responses to vaccination and safety profile of SerBut

Mice were orally gavaged with PBS, SerBut (twice daily, 24 mg/dose), or FTY720 (once daily, 0.02 mg/dose) starting on day -3 until the end of the experiment. On day 0, mice were immunized subcutaneously in the front hocks with 10 μg endotoxin-free ovalbumin (OVA), 50 μg alum, and 5 μg MPLA as described previously^60^. Mice were bled on day 9 and day 13, and plasma was analyzed for anti-OVA total IgG antibodies using a mouse anti-OVA IgG antibody assay kit (Chondrex). On day 13, mice were sacrificed, the hock-draining LNs and spleen were harvested, and cells were isolated for the immunostaining followed by flow cytometry analysis. One million cells from each spleen were seeded in a 96-well plate and incubated with OVA at 100 μg/mL for a 3-day, *ex vivo* restimulation. The supernatant of cell culture was collected, and cytokines were measured using a LEGENDplex™ mouse Th cytokine assay (BioLegend).

In addition, plasma samples collected on day 13 were analyzed for various markers of organ toxicity using a biochemistry analyzer (Alfa Wassermann Diagnostic Technologies) according to the manufacturer’s instructions. The panel included albumin, alanine amino-transferase, amylase, aspartate amino-transferase, blood urea nitrogen, calcium, creatine kinase, creatine, total bilirubin, and total protein.

### Statistical analysis

Statistical analysis and plotting of data was performed using Prism 9.0 (Graphpad), as indicated in the figure legends. One-way ANOVA with Dunnett’s,Tukey’s or Kruskal-Wallis test (if not normally distributed) for multiple comparisons was used in **Fig. 1d-h**, **Fig. 2b-h**, **Fig. 4b, d-n,** and **Fig. 6b, c, e, g, i**, **Extended Fig. 1b-g**, and **Extended Fig. 5**. Student’s t-test was used in **Fig. 3b-d, h-m**, **Fig. 5b, d, f-o**, and **Extended Fig. 2, 3.** Two-way ANOVA with Tukey’s or Bonferroni’s post-test was used in **Extended Fig. 4**. In **Fig. 3b**, **Fig. 4b**, **Fig. 5b**, and **Extended Fig. 1b**, the area under curve (AUC) values of clinical scores were compared using one-way ANOVA with Dunnett’s post-test, or Student’s t test. In **Fig. 4c** and **Fig. 5c**, the probability curve of EAE clinical scores remaining below 1.0 were compared between each two groups using the Log-rank (Mantel-Cox) test. Data represent mean ± s.e.m.; *n* is stated in the figure legend.

### Reporting Summary

Further information on research design is available in the Nature Research Reporting Summary linked to this article.

## Data availability

The main data supporting the results in this study are available within the paper and its Supplementary Information. Source data for the figures will be provided with this paper. All data generated in this study will be available from figshare.

## Acknowledgements

This work was supported by the Chicago Immunoengineering Innovation Center of the University of Chicago and the Alper Family Fund. We thank Yue Wang from Prof. Melody Swartz’s laboratory for providing mouse BMDCs. We thank Edward Ionescu from Prof. Cathryn Nagler’s laboratory for assisting with lamina propria isolation. We thank Dr. Abbey Lauterbach for helping on interpreting histology images. We thank the Cytometry and Antibody Technology Core Facility (Cancer Center Support Grant P30CA014599), the Human Tissue Resource Center, the Integrated Light Microscopy Core, and the Mass Spectrometry Facility (National Science Foundation instrumentation grant CHE-1048528) at the University of Chicago.

## Author contributions

J.A.H. oversaw all research. S.C., E.B., and J.A.H. designed most of experiments. S.C., M.M.R., T.N.B., A.J.S. synthesized materials. S.C., E.B., M.M.R., A.S., M.N.,T.N.B., J.W.R., K.H., P.A., L.A.H., A.T.A., K.C.R., L.S.S., I.P., R.P.W., A.D., and E.A.W. performed experiments. S.C. and E.B. analyzed experiments. S.C., E.B., and J.A.H. wrote the manuscript. All authors contributed to the article and approved the submitted version.

---

**Extended Fig. 1.**
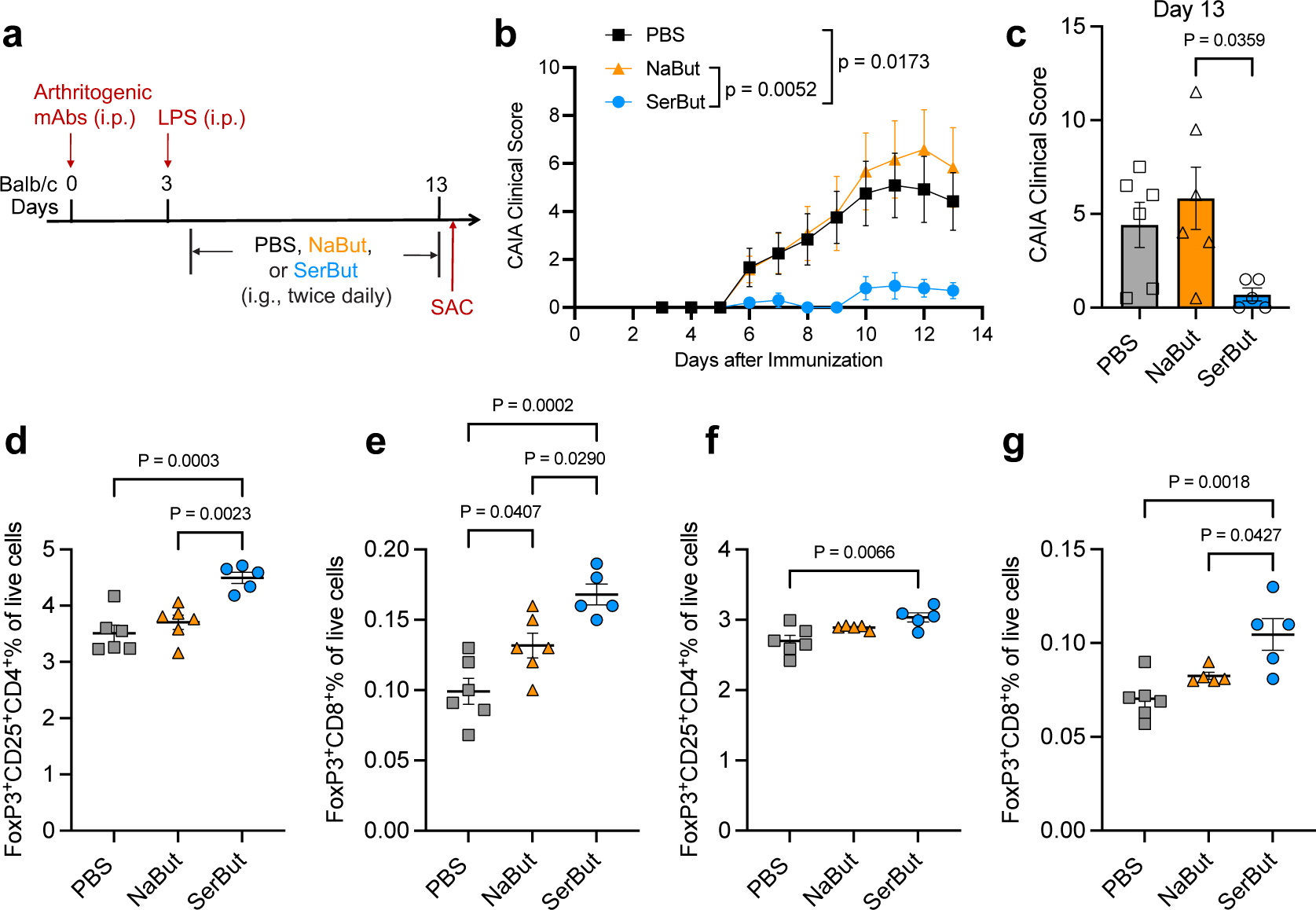
SerBut, but not NaBut, suppresses arthritis development. **a.** Experimental schema of the CAIA model. CAIA was induced by passive immunization with anti-collagen antibody cocktails on day 0, followed by intraperitoneal injection of lipopolysaccharide (LPS) on day 3. Starting on day 4, mice were orally gavaged with PBS (*n* = 6), NaBut (15 mg, molar equivalent to SerBut, *n* = 6) or SerBut (25 mg, *n* = 5) twice daily starting on day 4 until the end of the experiment. **B.** Arthritis scores in mice measured daily after the immunization. **C.** Arthritis scores from PBS or SerBut treated mice on day 13. **D, e.** Percentage of Foxp3^+^CD25^+^ regulatory CD4^+^ T cells (d) or Foxp3^+^ regulatory CD8^+^ T cells (e) of live cells in the hock-draining LNs. **f, g.** Percentage of Foxp3^+^CD25^+^ regulatory CD4^+^ T cells (f) or Foxp3^+^ regulatory CD8^+^ T cells (g) of live cells in the spleen. The experiments comparing PBS and SerBut were repeated three times and the results were consistent. NaBut were tested once as added in this experiment. Data represent mean ± s.e.m. Statistical analyses were performed using one-way ANOVA with Tukey’s post hoc test. P values less than 0.05 were shown.

**Extended Fig. 2.**
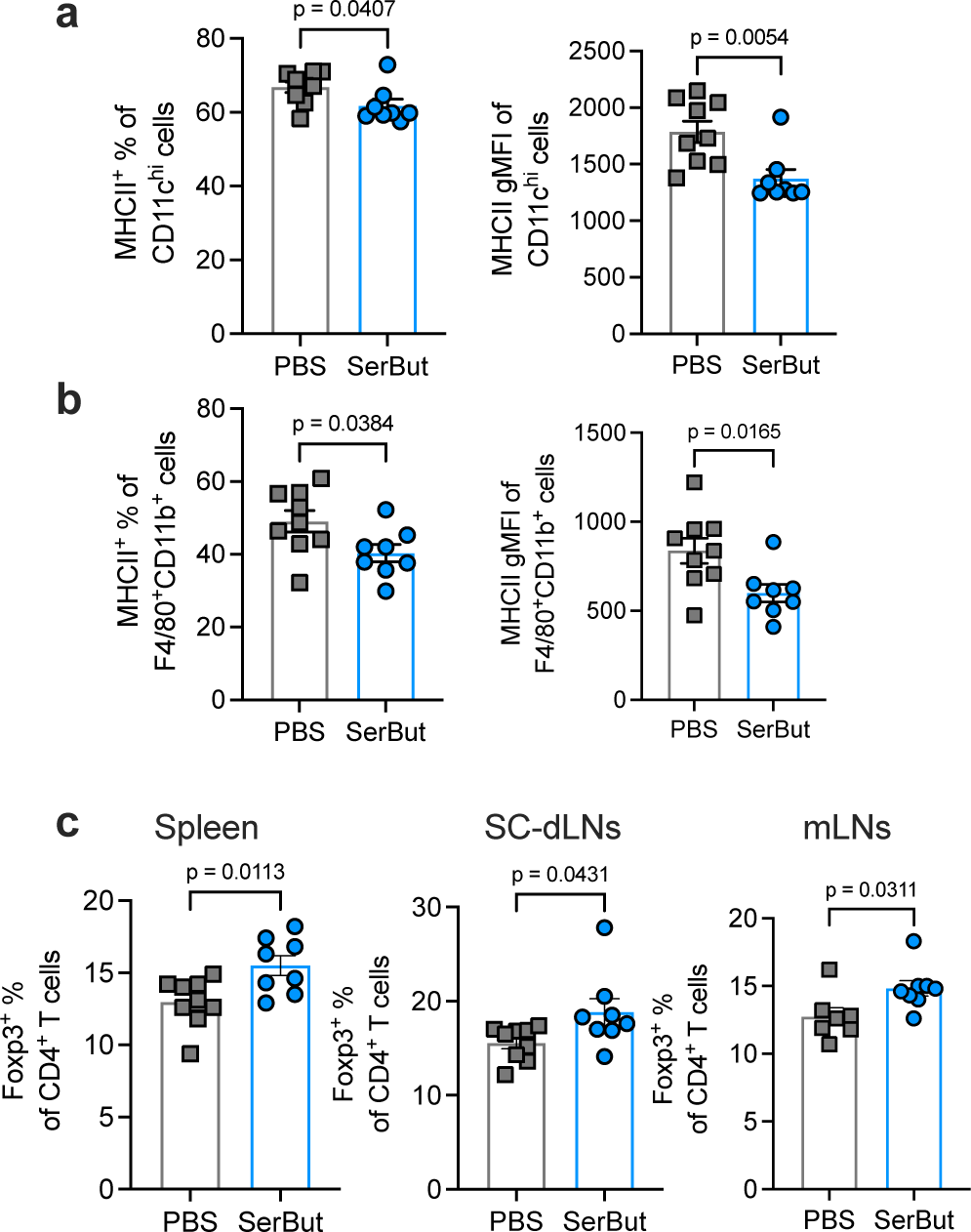
The immunological effects in peripheral tissues from SerBut treatment the EAE model from the experiment in **Figure 5**. **a, b.** The percentage of MHC class II^+^% and geometric mean fluorescent intensity (gMFI) of CD11c^hi^ dendritic cells (**a**) or F4/80+CD11b+ macrophages (**b**) isolated from mesenteric LN. **c.** The percentage of Foxp3^+^ of CD4^+^ T cells in the spleen, spinal cord-draining LNs (SC-dLNs), or mesenteric LNs (mLNs). Data represent mean ± s.e.m. Statistical analyses were performed using Student’s t-test.

**Extended Fig. 3.**
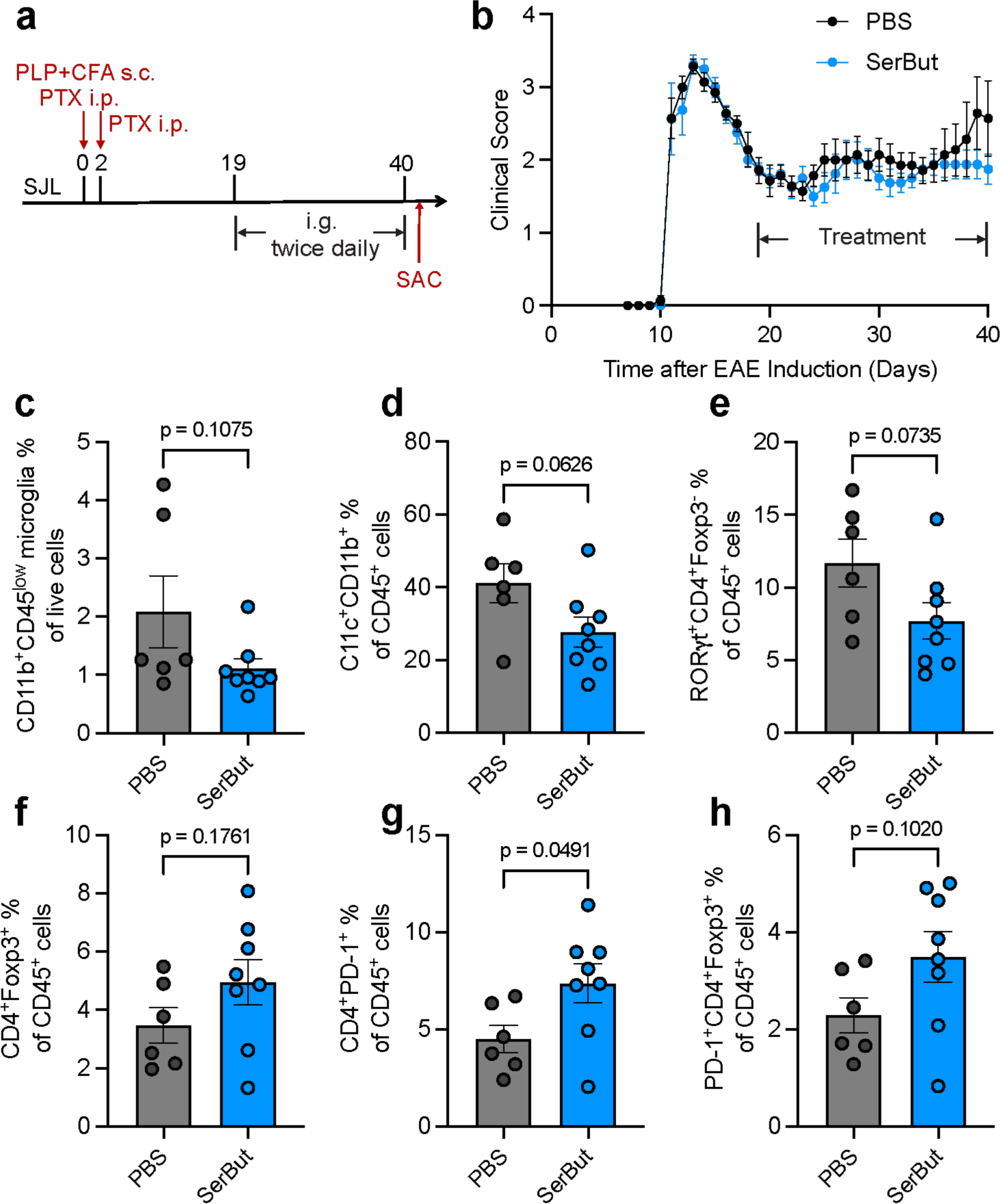
SerBut exhibits moderate immunomodulation in the spinal cord following therapeutic treatment in relapsing-remitting EAE. **a.** Experimental schema. EAE was induced in SJL/J mice using PLP_139-151_/CFA with 100 ng pertussis toxin. Starting on day 19, mice were regrouped into two treatment groups with equivalent average clinical score, and received twice daily oral gavage of either PBS (*n* = 6) or SerBut (*n* = 8) at 24 mg/dose. **b.** Disease progression as indicated by the clinical score. **c.** The percentage of CD11b^+^CD45^low^ microglia cells of live cells in the spinal cord. **d-h.** The percentage of CD11b^+^CD11c^+^(d), RORγ^+^CD4^+^Foxp3^-^(e), CD4^+^Foxp3^+^(f), CD4^+^PD-1^+^(g), PD-1^+^CD4^+^Foxp3^+^(h) of CD45^+^ cells in the spinal cord. Data represent mean ± s.e.m. Statistical analyses were performed using Student’s t-test.

**Extended Fig. 4.**
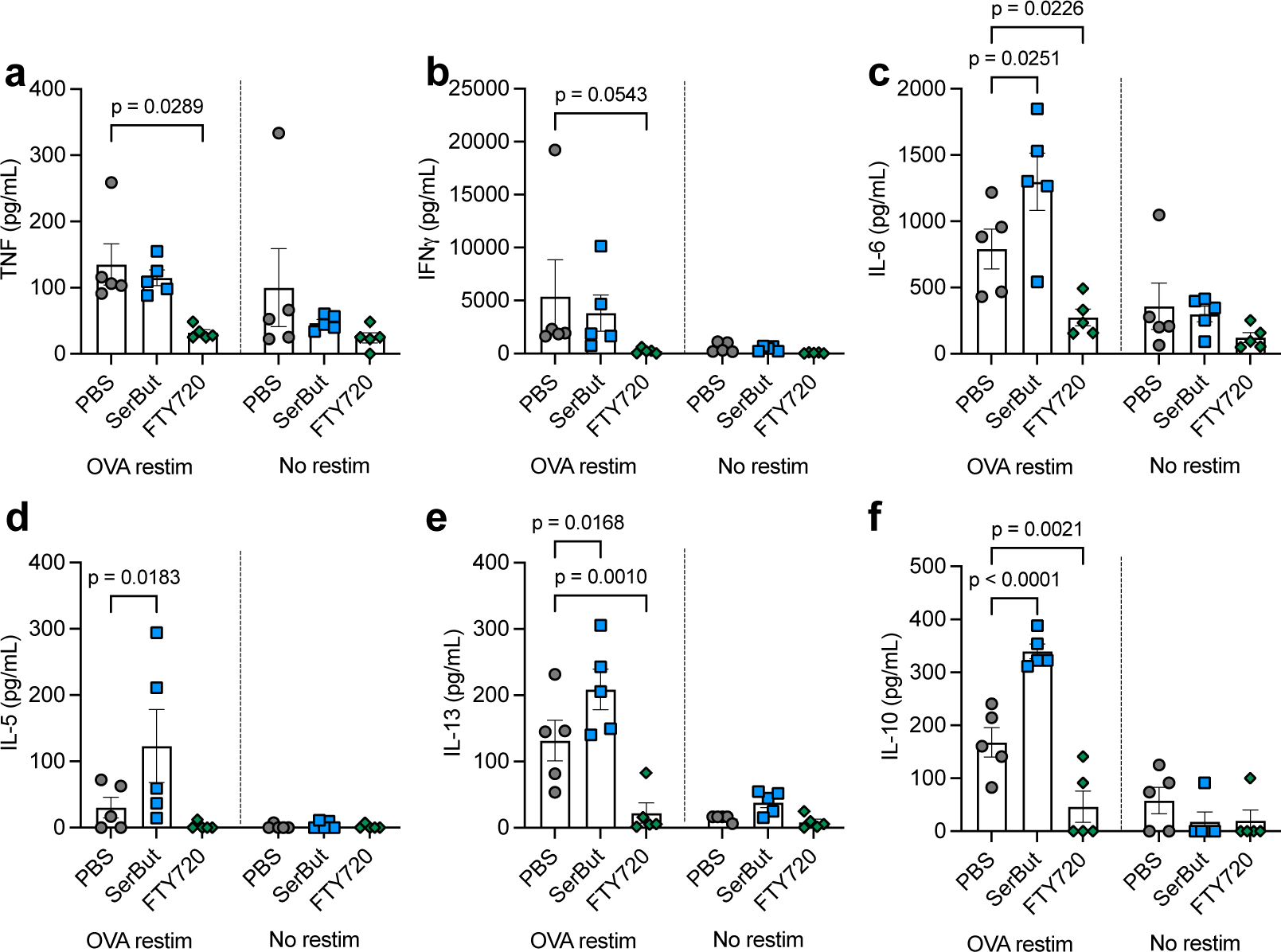
SerBut’s impact on immune responses to vaccination in comparison to FTY720. Production of cytokines TNFa (a), IFNγ (b), IL-6 (c), IL-5 (d), IL-13 (e), and IL-10 (f), measured in the supernatant of isolated splenocytes upon restimulation with OVA protein for 3 days. *n* = 5 mice per group. Data represent mean ± s.e.m. Statistical analyses were performed using two-way ANOVA with Dunnett’s post hoc test. P values less than 0.05 were shown.

**Extended Fig. 5.**
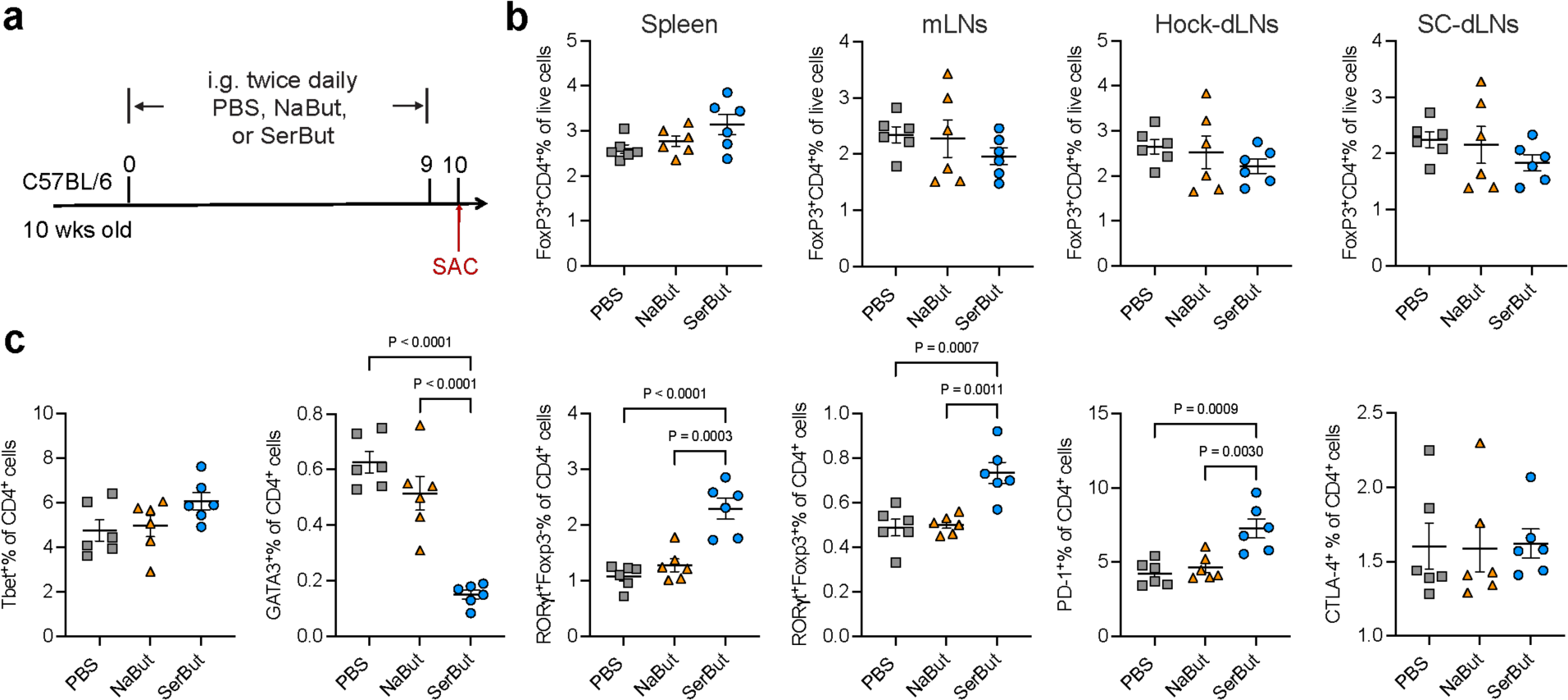
Immunological effects of SerBut treatment in healthy C57BL/6 mice on the T cells in the spleen or lamina propria. **a.** Experimental schema. C57BL/6 mice at 10 wks old were oral administered with PBS (*n* = 5), NaBut (15 mg, molar equivalent to SerBut, *n* = 6) or SerBut (25 mg, *n* = 6), twice daily for 10 days, and sacrificed for cellular analysis. **b**. CD4^+^Foxp3^+^ Tregs percentage of live cells in spleen, mesenteric LNs (mLNs), hock-draining LNs (hock-dLNs), and spinal cord-draining LNs (SC-dLNs). **c**. Tbet^+^, GATA3^+^, RORγ^+^ Foxp3^-^, RORγ^+^ Foxp3^+^, PD-1^+^, CTLA-4^+^ of CD4^+^ T cells in the spleen. Data represent mean ± s.e.m. Statistical analyses were performed using one-way ANOVA with Tukey’s post hoc test. P values less than 0.05 were shown.

