## Supplementary Information for "Suppression of autoimmune arthritis and neuroinflammation via an amino acid-conjugated butyrate prodrug with enhanced oral bioavailability"

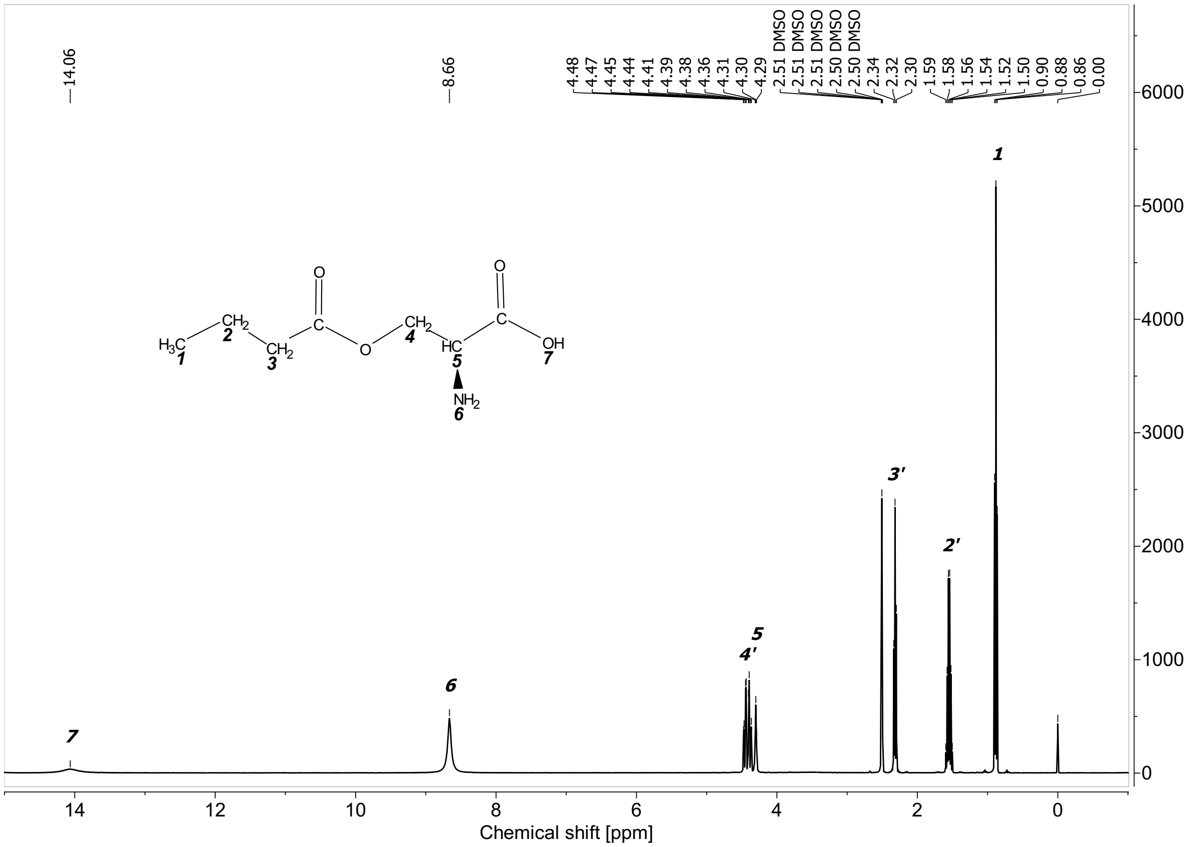

**Figure S1.** $^1$H NMR spectrum of *O*-butyryl-*L*-serine (SerBut) (500MHz, DMSO-d$_6$) [ppm]: 0.88 (3H, t), 1.55 (2H, m), 2.32 (2H, t), 4.30 (1H, t), 4.43 (2H, d), 8.66 (2H, s), 14.06 (1H, s).

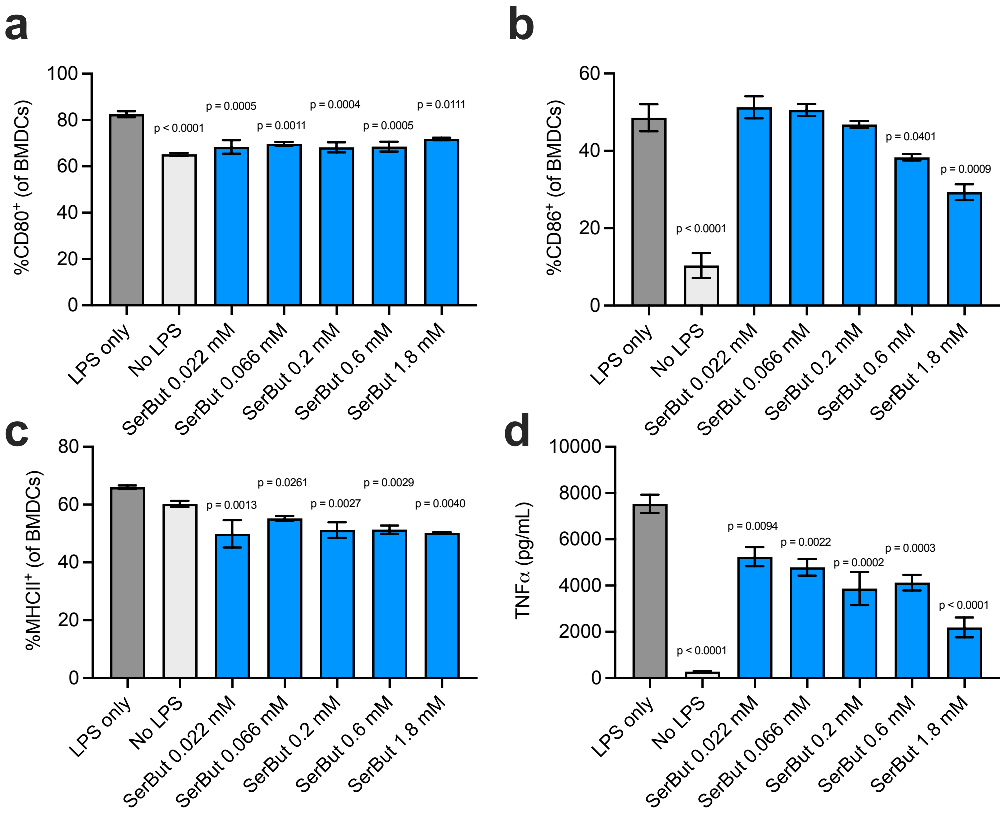


**Figure S2.** SerBut suppressed BMDC activation. BMDCs incubated with SerBut at a series of concentrations for 24 hr, followed by LPS stimulation for 18 hr. **a-c**, Percentage of $CD80^+$, $CD86^+$, or $MHCII^+$ cells of BMDCs analyzed by flow cytometry. **d**. TNFα concentration in the cell culture supernatant of BMDCs. Data represent mean ± s.e.m. Statistical analyses were compared between LPS only group with no LPS or SerBut-treated group, performed using a one-way ANOVA with Dunnett's test. P value less than 0.05 were shown.

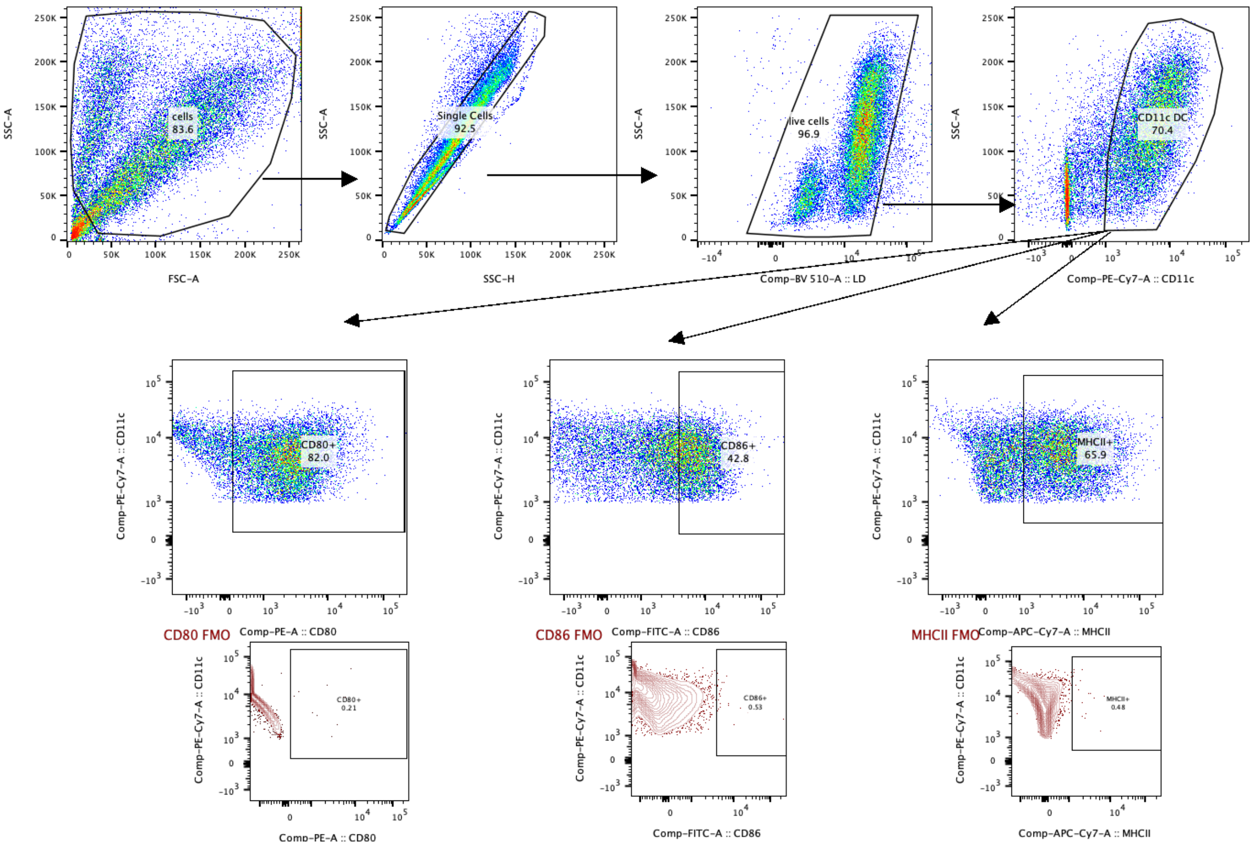


**Figure S3.** Flow cytometry gating strategy for the identification of MHCII$^{+}$, CD80$^{+}$, or CD86$^{+}$ cells of BMDCs. Representative FMOs (fluorescence minus one) were included for CD80, CD86, and MHCII gating strategy. This gating strategy was used in Fig.1d-f, and Fig. S2.

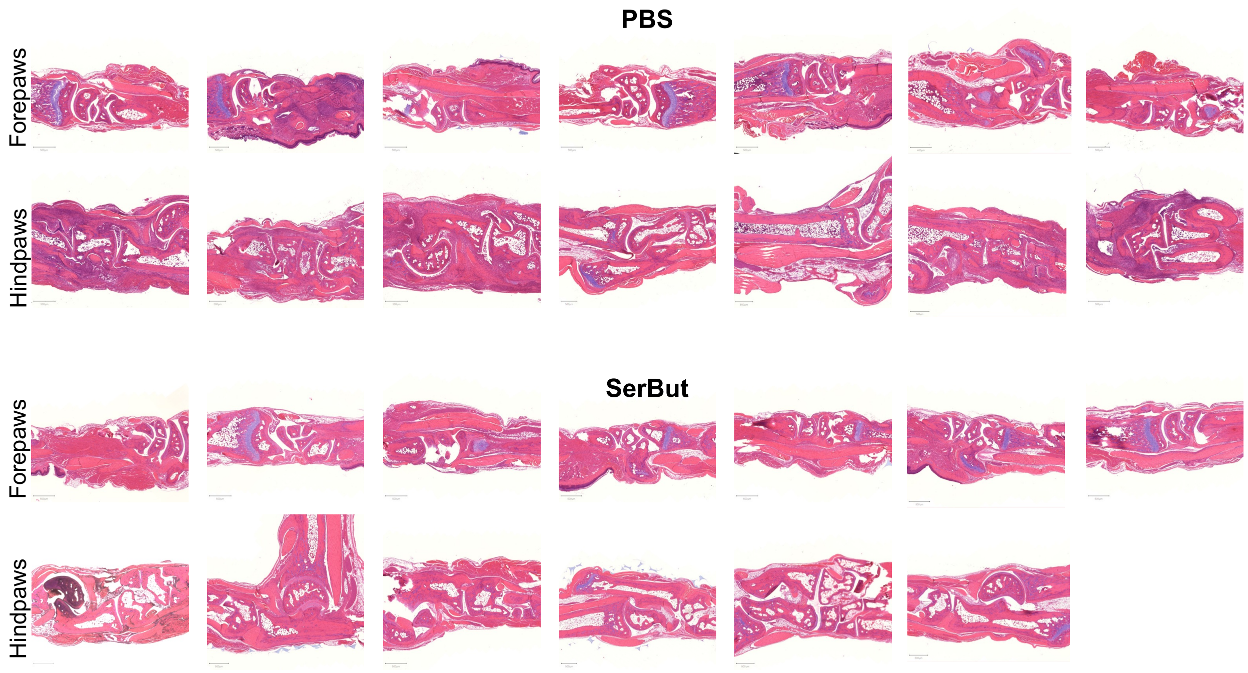


**Figure S4.** Histology images of mouse joints from paws stained with hematoxylin and eosin, from the experiment in Fig.2. (One hindpaw sample from SerBut group was lost during processing.)

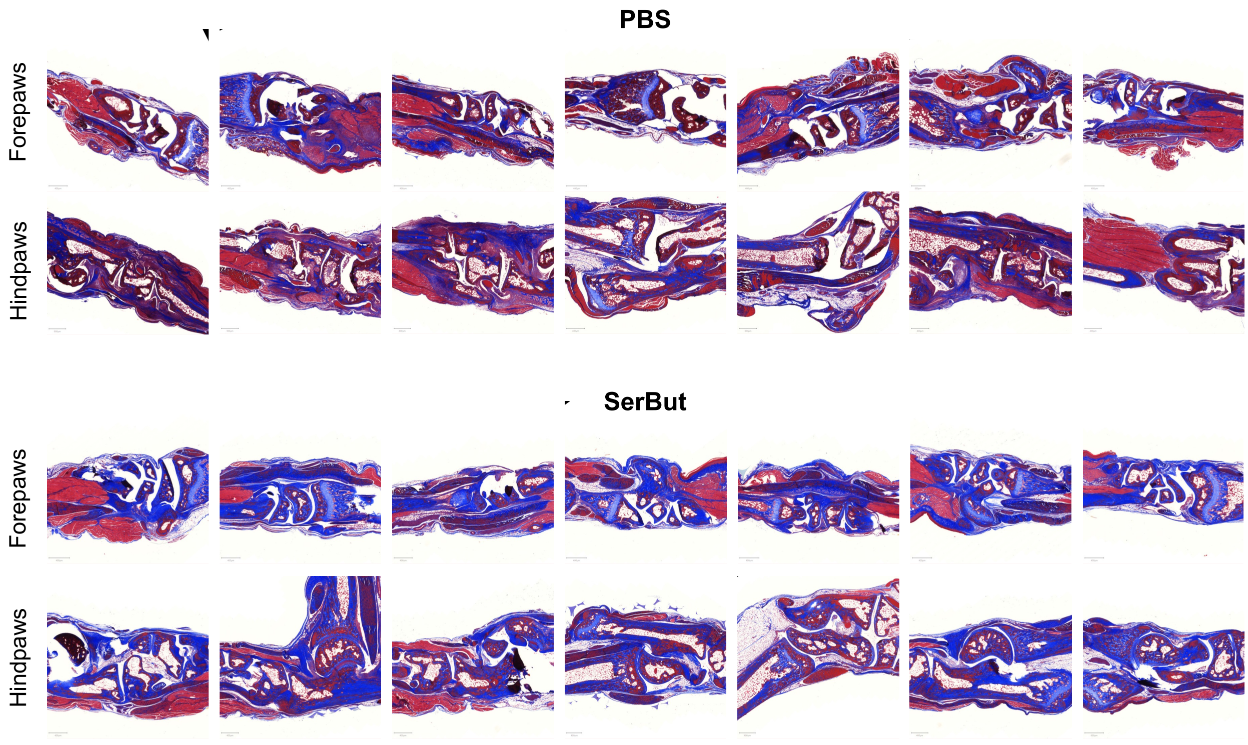


**Figure S5.** Histology images of mouse joints from paws stained with Masson's trichrome, from the experiment in Fig. 2. Blue represents collagen staining.

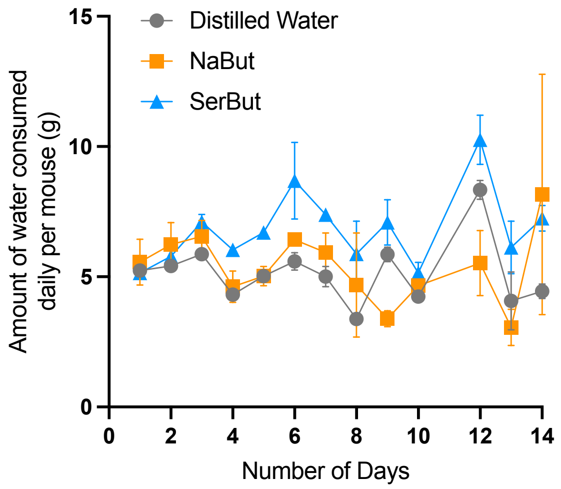


**Figure S6.** Measurement of the amount of drinking water consumed by healthy, C57BL/6 mice. Distilled water alone or supplemented with 100mM NaBut or SerBut was administered to the mice over a 14-day period.

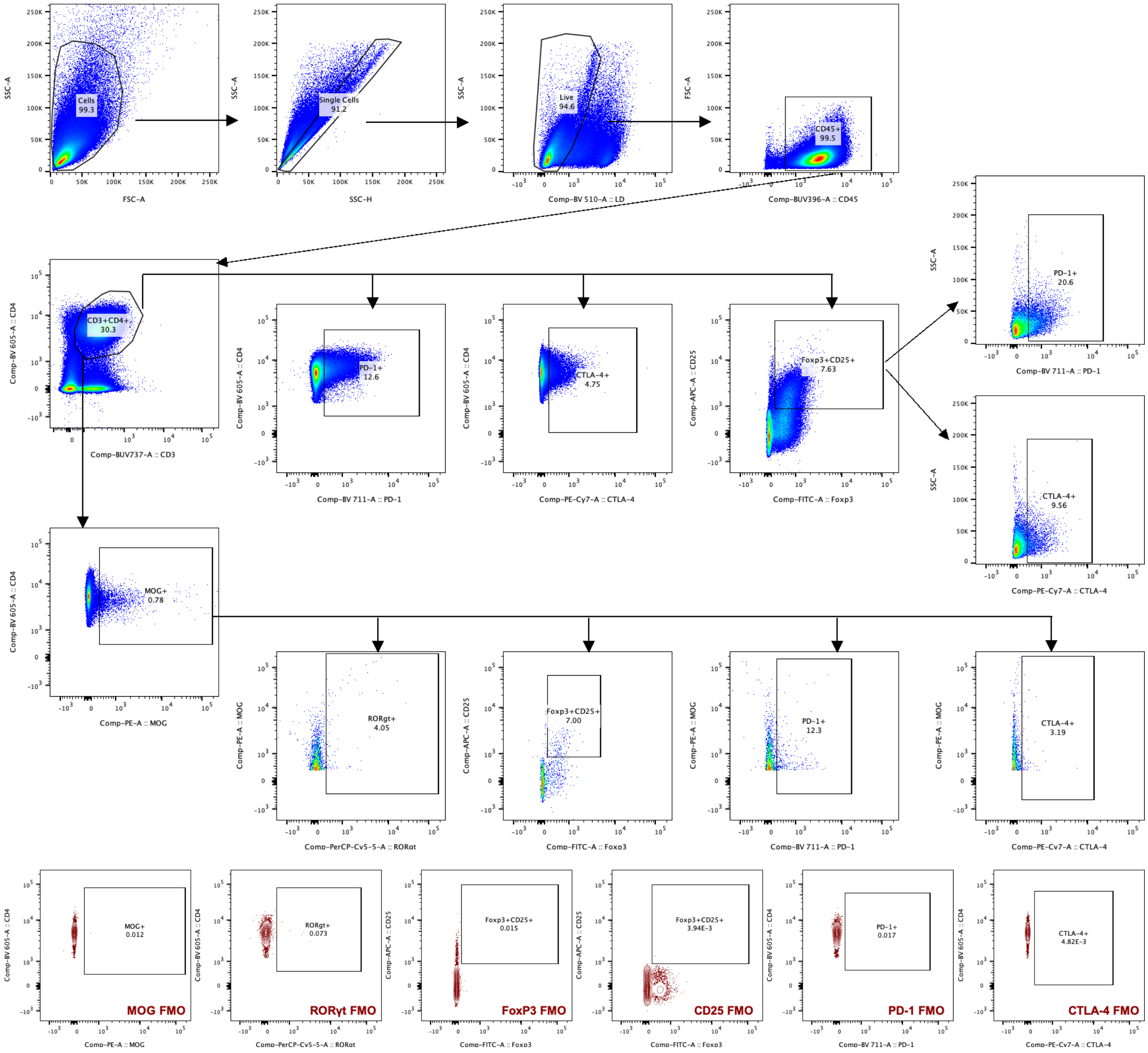


**Figure S7.** Flow cytometry gating strategy for the identification of PD-1$^+$, CTLA-4$^+$, or Foxp3$^+$CD25$^+$ of CD4$^+$ T cells, PD-1$^+$ or CTLA-4$^+$ of Foxp3$^+$CD25$^+$CD4$^+$ Tregs, MOG tetramer-positive CD4$^+$ or CD4$^+$RORγt$^+$ T cells, and Foxp3$^+$CD25$^+$, PD-1$^+$, CTLA-4$^+$ of MOG tetramer-positive CD4$^+$ T cells. This gating strategy was used in Fig. 4, 5.

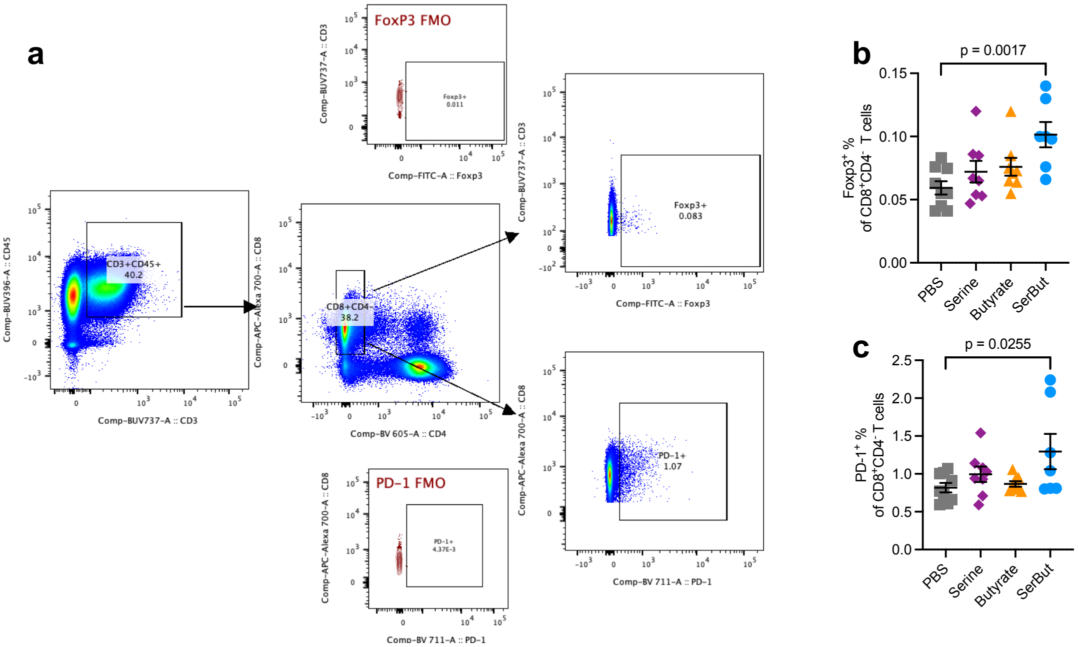


**Figure S8.** The gating strategy (**a**) and the percentage of Foxp3$^+$ (**b**) or PD-1$^+$ (**c**) of CD8$^+$CD4$^-$ T cells in the spinal cord-draining lymph nodes (SC-dLNs, iliac and cervical LNs) measured by flow cytometry, from the experiment in Fig. 4. Data represent mean ± s.e.m. Statistical analyses were compared between PBS and each treatment group using one-way ANOVA with Dunnett's test. P values less than 0.05 were shown.

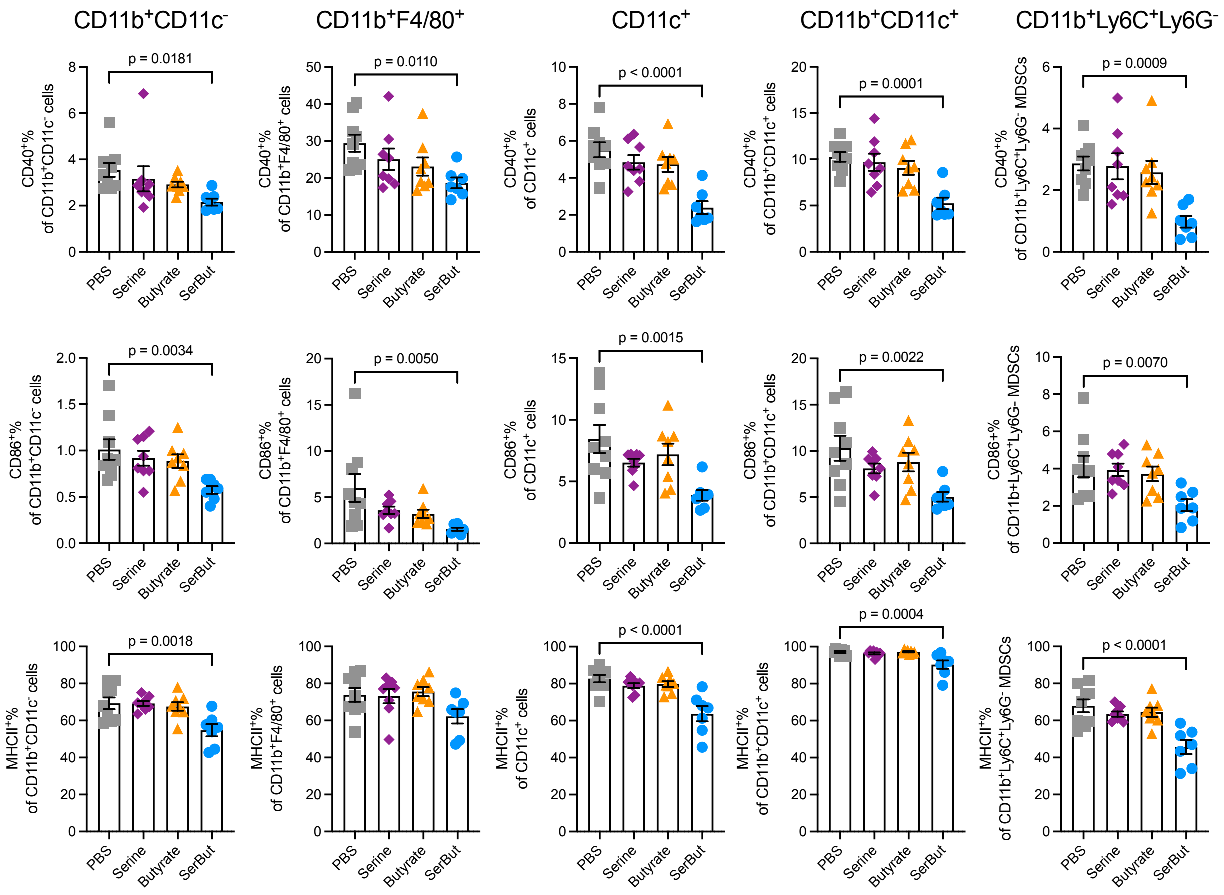


**Figure S9.** The percentage of co-stimulatory molecule (CD40$^{+}$ and CD86$^{+}$) and MHCII$^{+}$ cells of myeloid cells in the SC-dLNs from Fig. 4n. Data represent mean ± s.e.m. Statistical analyses were compared between PBS and each treatment group using one-way ANOVA with Dunnett's test. P values less than 0.05 were shown.

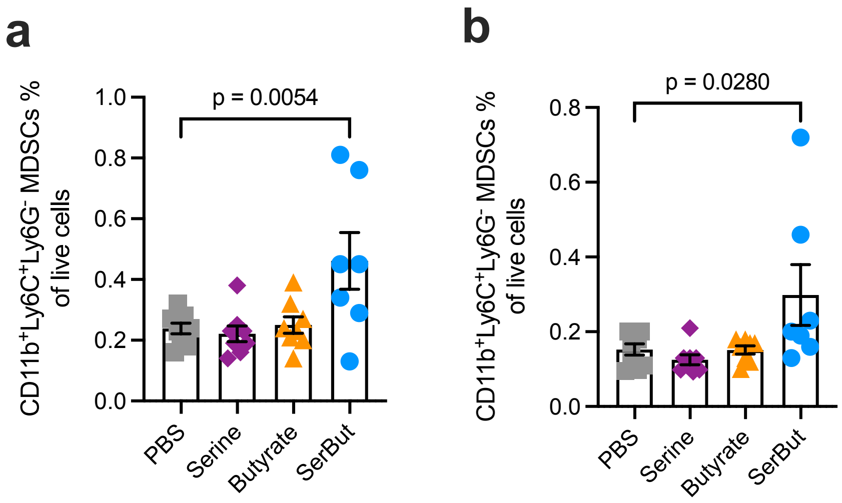


**Figure S10.** The percentage of $CD11b^{+}Ly6C^{+}Ly6G^{-}$ cells in the spinal cord-draining LNs (**a**) or mesenteric LNs (**b**) from the experiment in Fig. 3. Data represent mean ± s.e.m. Statistical analyses were compared between PBS and each treatment group using one-way ANOVA with Dunnett's test. P values less than 0.05 were shown.

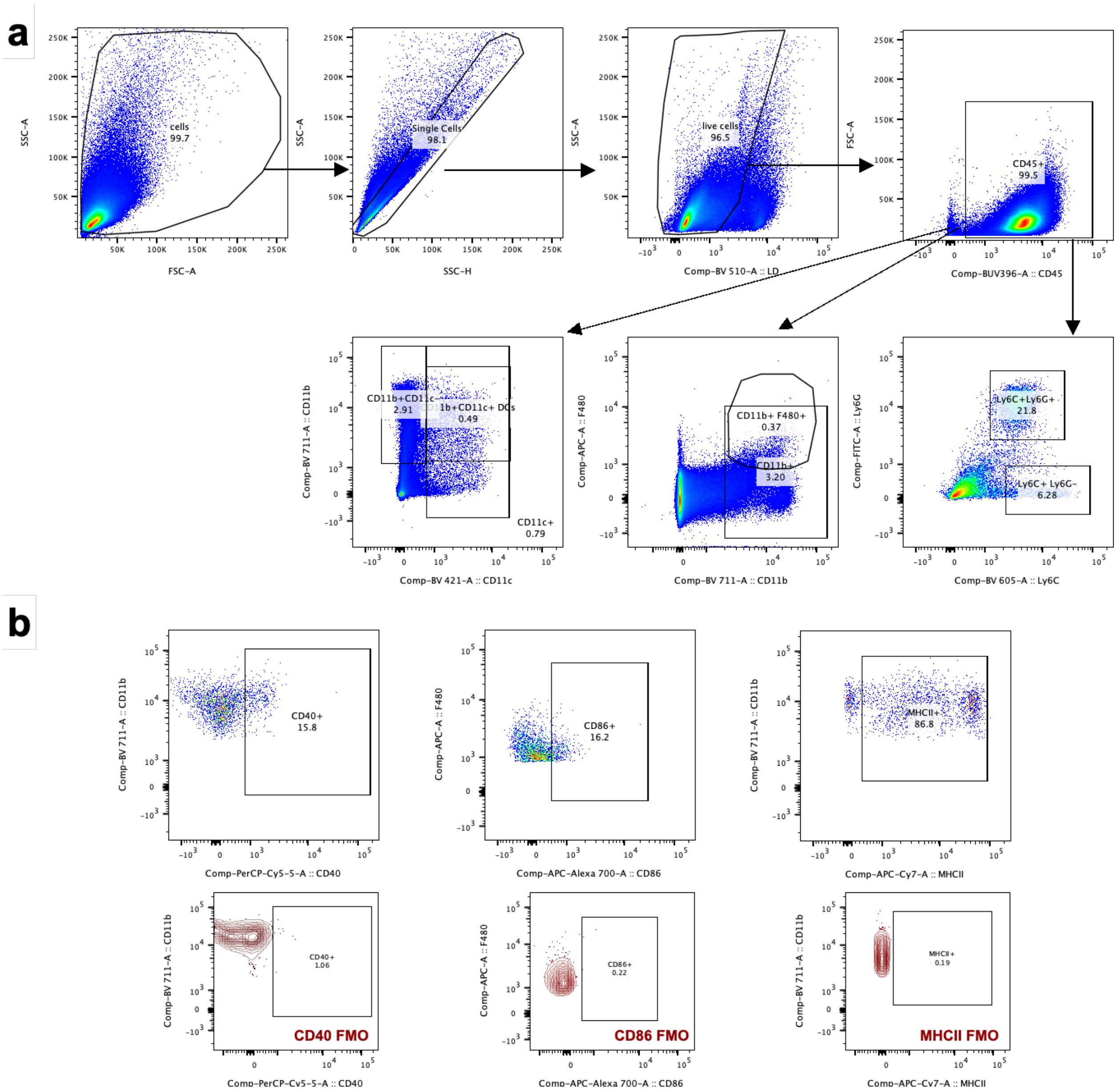


**Figure S11. a.** Flow cytometry gating strategy for CD11b$^{+}$CD11c$^{-}$, CD11b$^{+}$F4/80$^{+}$, CD11c$^{+}$, CD11b$^{+}$CD11c$^{+}$, CD11b$^{+}$Ly6C$^{+}$Ly6G$^{-}$ cells **b**. Representative gating strategy for CD40$^{+}$, CD86$^{+}$, or MHCII$^{+}$ cells of CD11b$^{+}$F4/80$^{+}$ with FMO. This gating strategy was used in Fig. 4n and Fig. S9, S10.

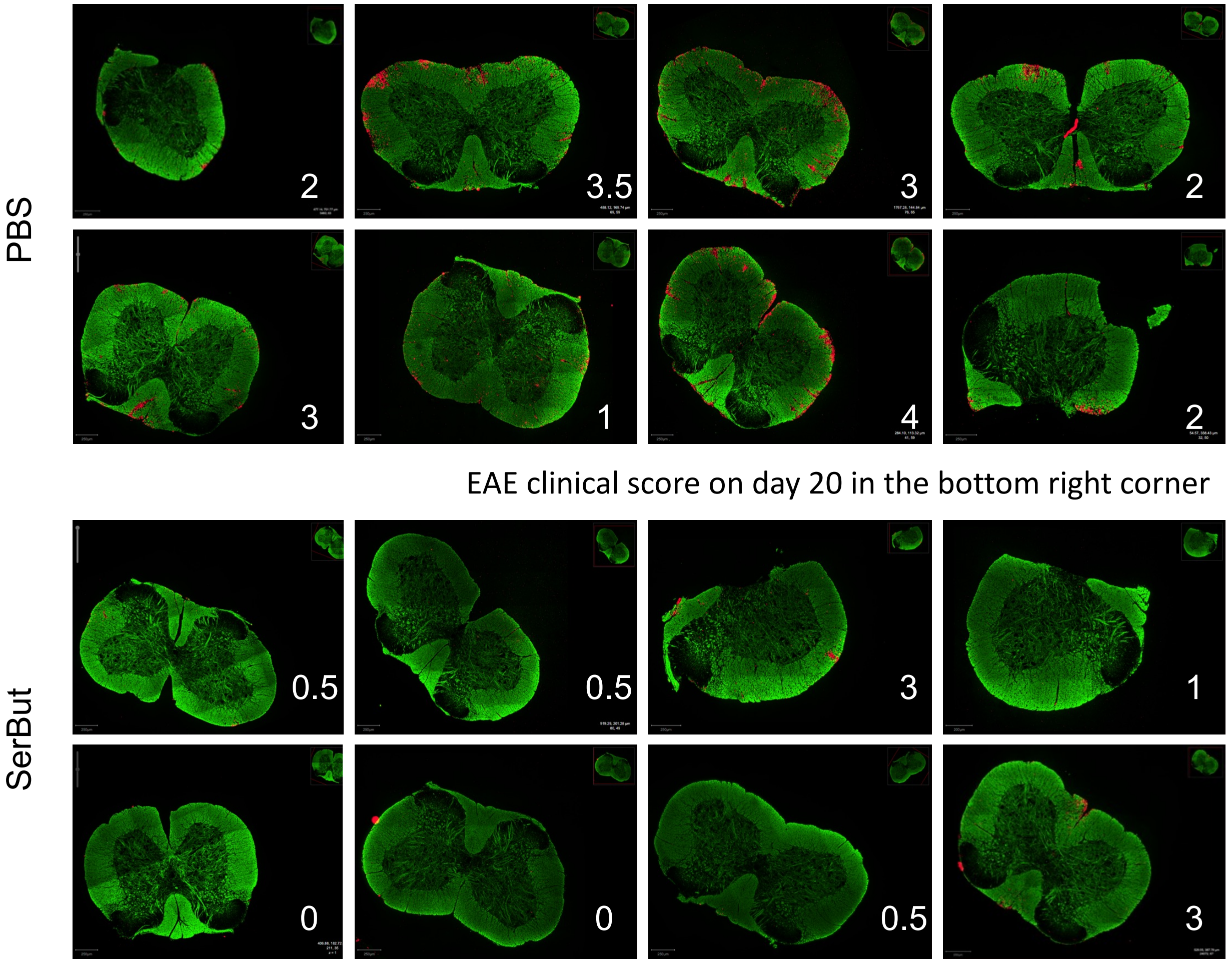


**Figure S12.** Immunofluorescent images of spinal cord sections from mice treated with PBS or SerBut, taken from the experiment in Fig. 4. The EAE clinical score on day 20 is displayed in the bottom right corner of each image. Red: anti-CD45 staining; green: anti-myelin basic protein (MBP) staining.

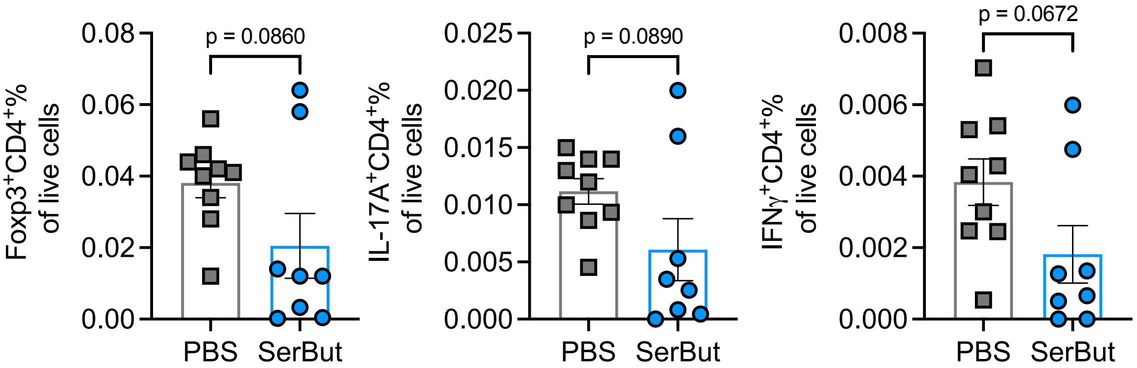


**Figure S13.** The percentage of $Foxp3^{+}CD4^{+}$, IL-17$A^{+}CD4^{+}$, and IFN$\gamma^{+}CD4^{+}$ cells of live cells in the spinal cord, from the experiment in Fig. 5. Data represent mean ± s.e.m. Statistical analyses were performed using Student's t-test.

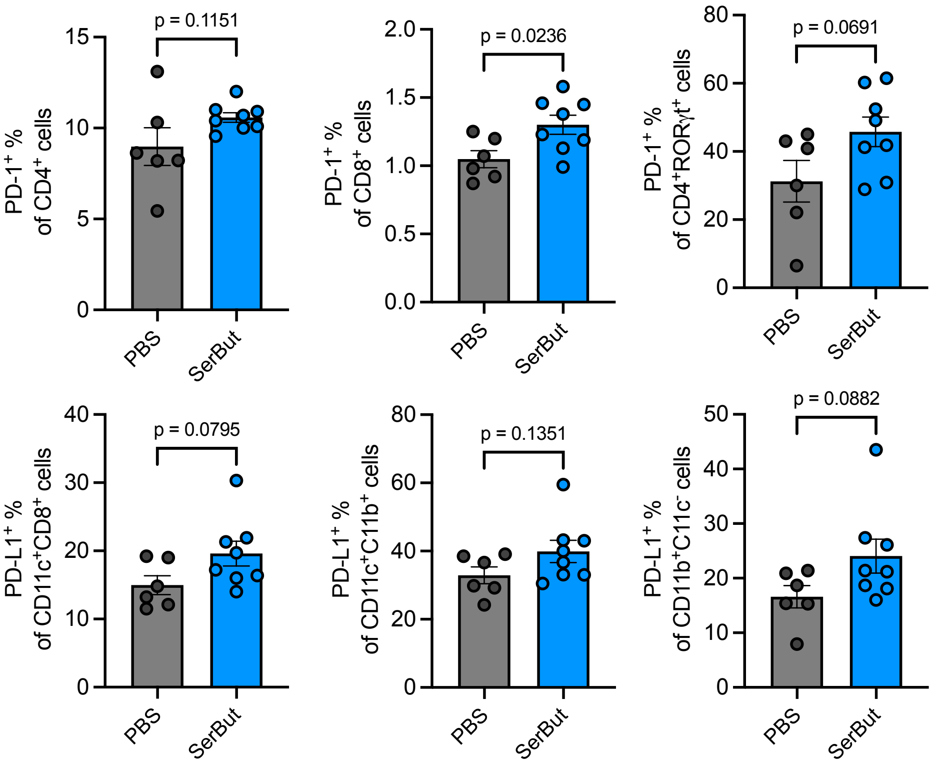


**Figure S14. The effect of SerBut on PD-1 expression of T cells, and PD-L1 expression on myeloid cells in mesenteric LNs from the therapeutic EAE experiment in Extended Fig. 3.** Data represent mean ± s.e.m. Statistical analyses were performed using Student's t-test.

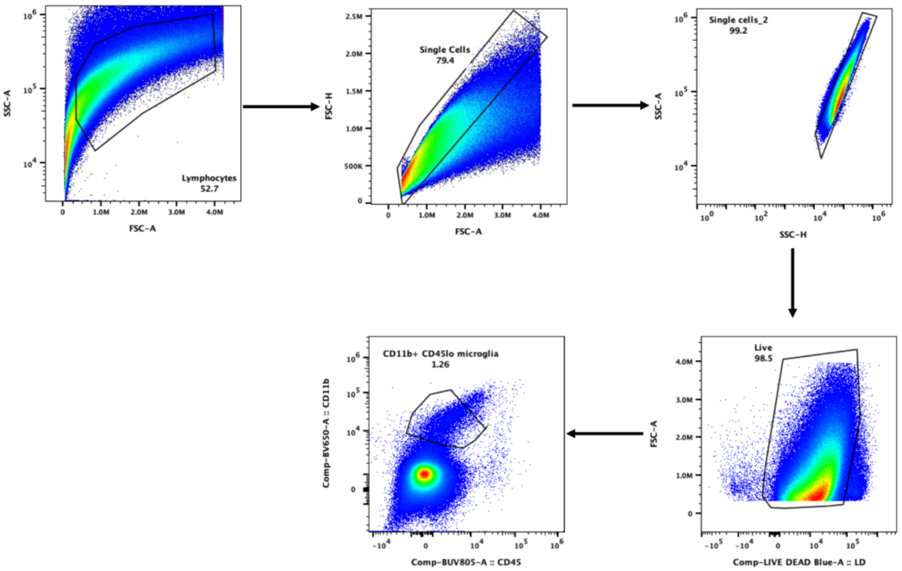


**Figure S15**. Flow cytometry gating strategy for CD11b$^+$CD45$^{low}$ spinal cord microglia in Extended Fig. 3c.

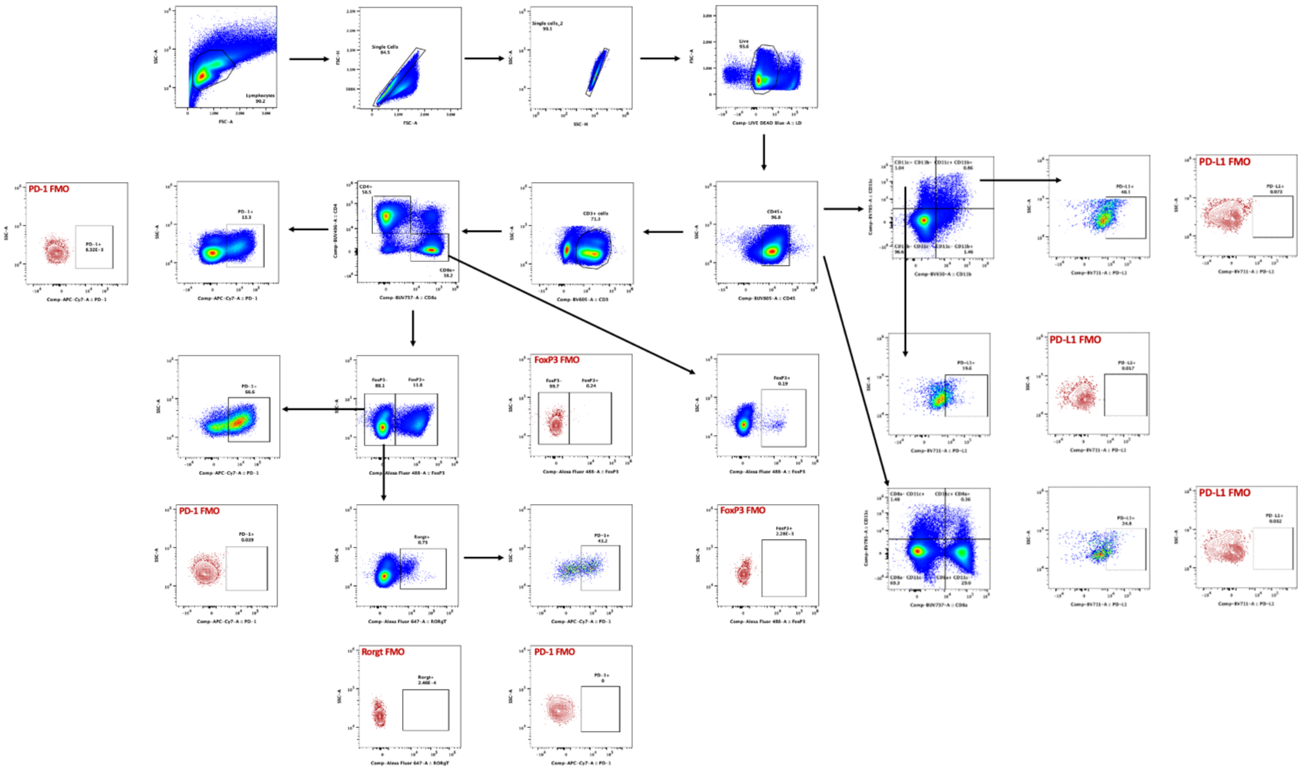


**Figure S16**. Flow cytometry gating strategy for RORγt$^+$ FoxP3$^-$ CD4$^+$ T cells, FoxP3+ CD4$^+$ T cells, PD-1$^+$ CD4$^+$ T cells, PD-1$^+$ FoxP3$^+$ CD4$^+$ T cells, PD-L1$^+$ CD11c$^+$ CD8$^+$ cells, CD11b$^+$ CD11c$^+$ myeloid cells, PD-L1$^+$ CD11c$^+$ CD11b$^+$ cells, and PD-L1$^+$ CD11c$^+$ CD11b$^-$ cells. This gating strategy was used in Extended Fig. 3, and Supplementary Fig. S14.

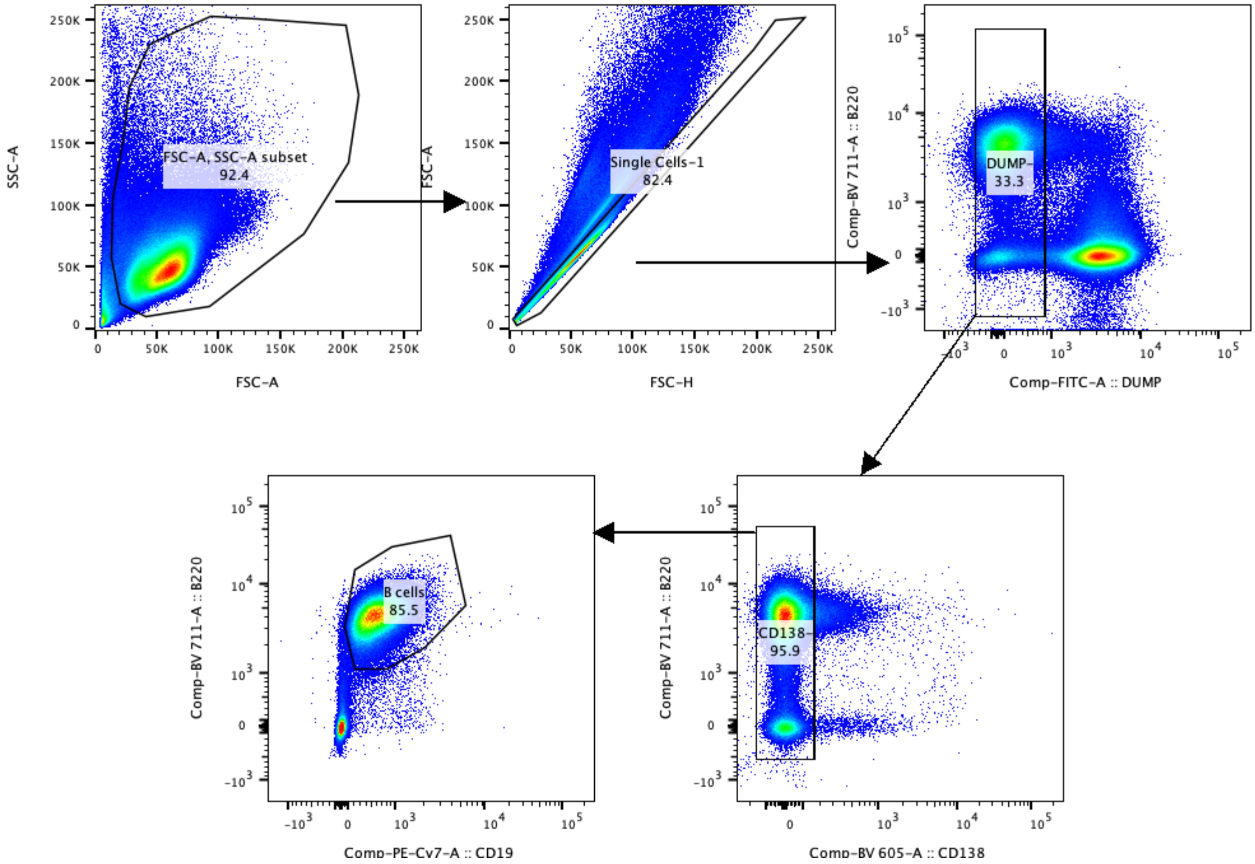


**Figure S17.** Flow cytometry gating strategy for CD19$^{+}$B220$^{+}$, B cells in Fig. 6d, e. Dump gate to exclude cells stained with FITC antibody against F4/80, CD11c, Gr-1, CD4, and CD8.

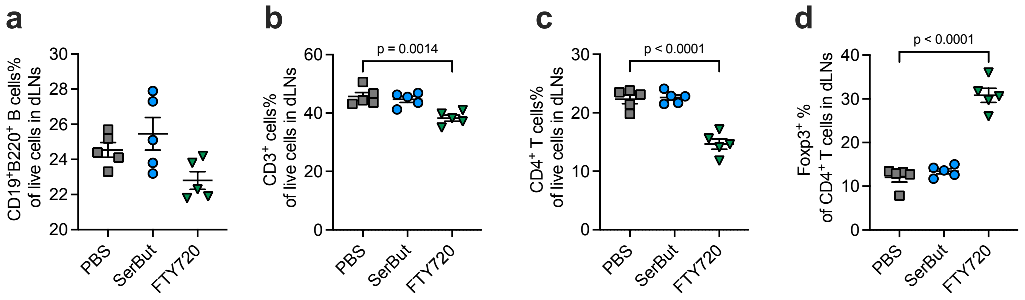


**Figure S18.** The percentages of CD19$^+$B220$^+$ (**a**), CD3$^+$ (**b**), CD4$^+$ (**c**) of total live cells and Foxp3$^+$ of CD4$^+$ T cells (**d**) in the hock-draining LNs, from the experiment in Fig. 6. Data represent mean ± s.e.m. Statistical analyses were performed using one-way ANOVA with Dunnett's post hoc test. P values less than 0.05 were shown.

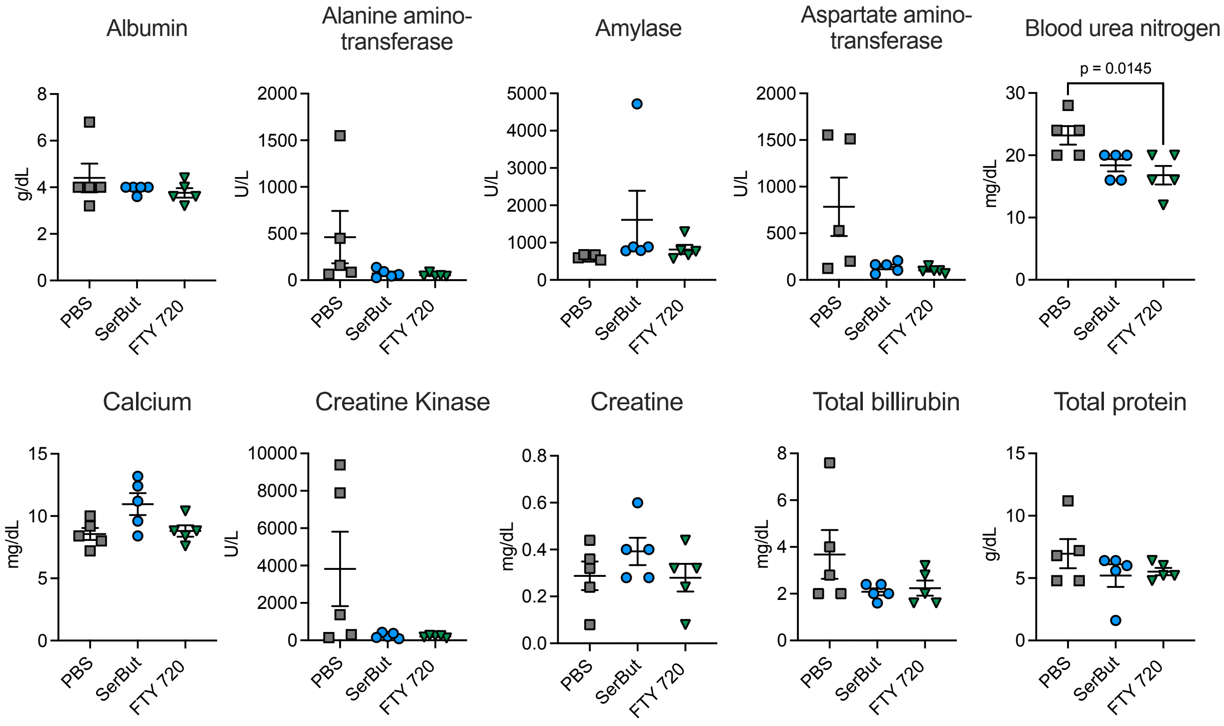


**Figure S19.** Serological toxicity analysis of mouse serum samples from the experiment in Figure 5. Data represent mean ± s.e.m. Statistical analyses were compared between every two groups using one-way ANOVA with Tukey's test. P values less than 0.05 were shown.

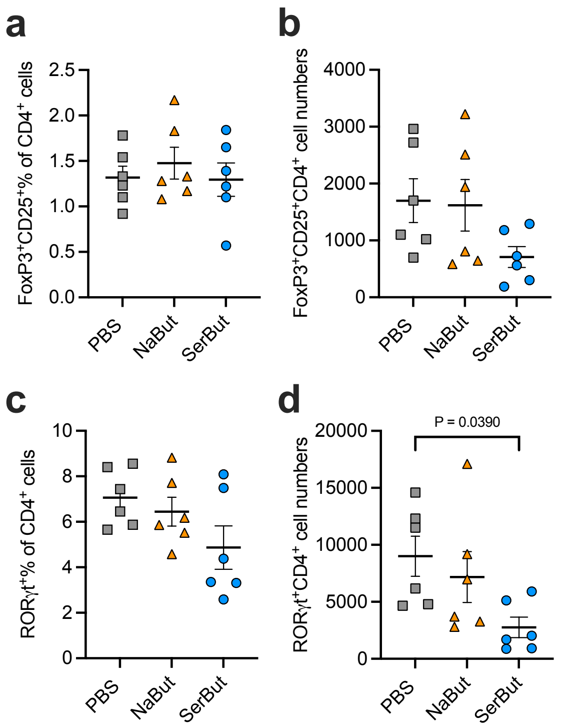


**Figure S20. Immunological effects of SerBut treatment in healthy C57BL/6 mice on the Tregs and Th17 cells in the ileum lamina propria from healthy mice treated with PBS, NaBut, or SerBut from Extended Fig. 5. a, b.** The percentage of Foxp3$^{+}$CD25$^{+}$ of CD4$^{+}$ T cells and their cell numbers in the lamina propria. **c, d.** The percentage of RORγt$^{+}$ of CD4$^{+}$ T cells and their cell numbers in the laminar propria. Data represent mean ± s.e.m. Statistical analyses were compared between PBS and each treatment group using one-way ANOVA with Dunnett's test. P values less than 0.05 were shown.

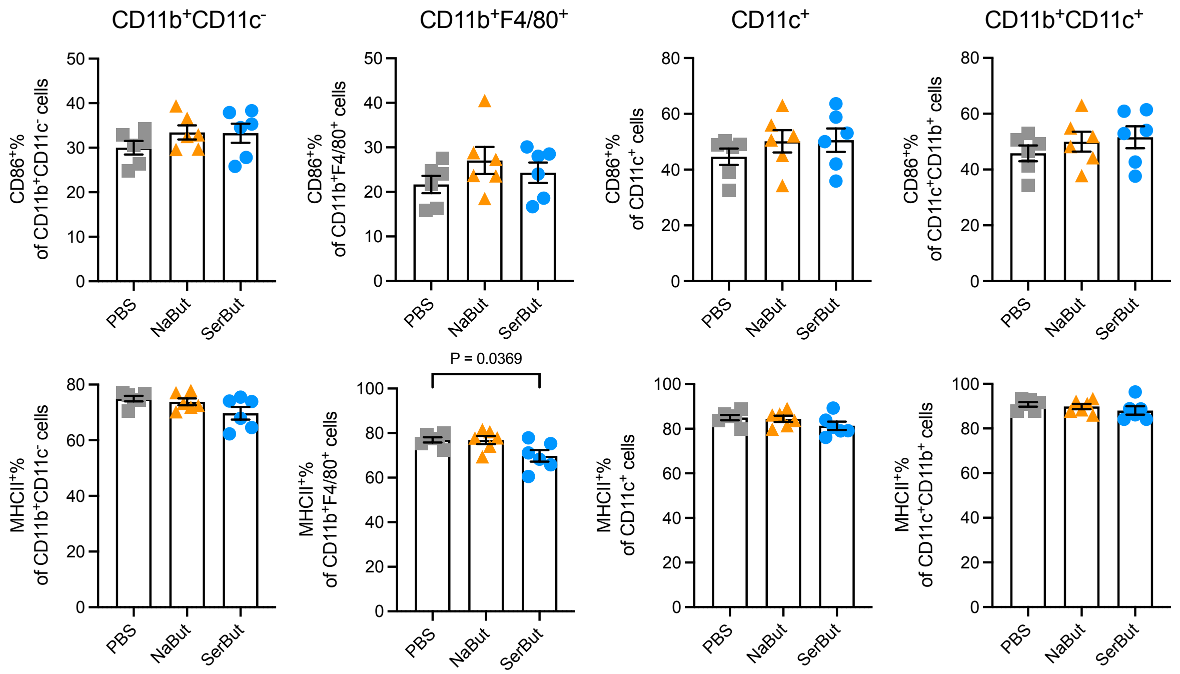


**Figure S21.** The percentage of co-stimulatory molecule CD86$^+$ and MHCII$^+$ cells of myeloid cells in the mesenteric LNs from healthy mice treated with PBS, NaBut, or SerBut from Extended Fig. 5. Data represent mean ± s.e.m. Statistical analyses were compared between PBS and each treatment group using one-way ANOVA with Dunnett's test. P values less than 0.05 were shown.

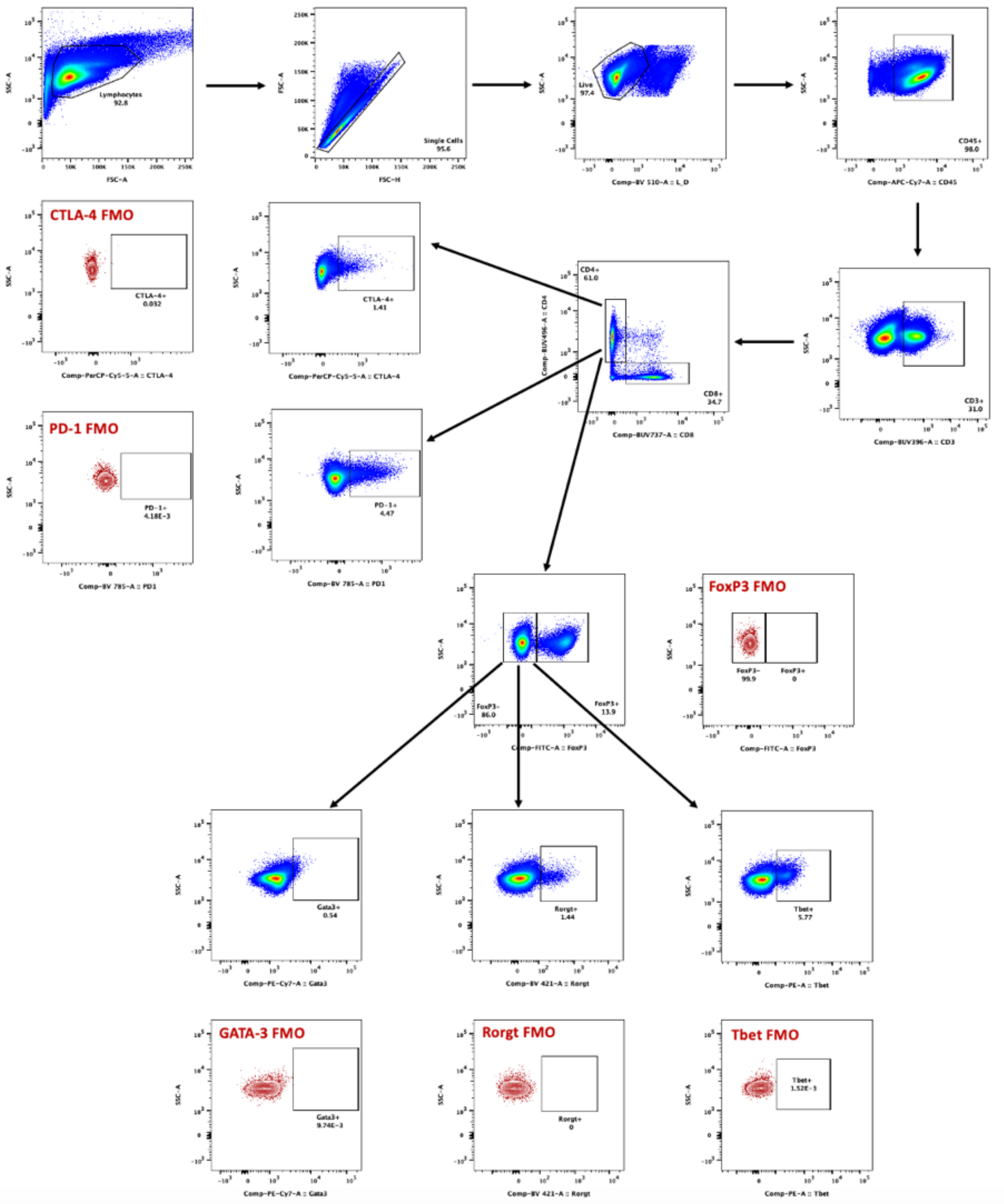


**Figure S22**. Flow cytometry gating strategy for PD-1$^{+}$ CD4$^{+}$ T cells, CTLA-4$^{+}$ CD4$^{+}$ T cells, RORγt$^{+}$ FoxP3$^{-}$ CD4$^{+}$ T cells, Gata3$^{+}$ FoxP3$^{-}$ CD4$^{+}$ T cells, and Tbet$^{+}$ FoxP3$^{-}$ CD4$^{+}$ T cells in the spleen in Extended Fig. 5c.
